# Nitric Oxide-Releasing Thixotropic Hydrogels as Antibacterial and Hemocompatible Catheter Locks

**DOI:** 10.1101/2025.08.06.668907

**Authors:** Wuwei Li, Loren Liebrecht, Surendra Poudel, Rebecca Goodhart, Sayaji More, Jade Montano, Derek Lust, Qingguo Xu, Martin Mangino, Xuewei Wang

**Affiliations:** Department of Chemistry, Virginia Commonwealth University, Richmond, Virginia 23284, United States; Department of Surgery, Virginia Commonwealth University, Richmond, Virginia 23223, United States; Department of Pharmaceutics, Virginia Commonwealth University, Richmond, Virginia 23298, United States

**Keywords:** Catheter, infection, hydrogel, nitric oxide, lock therapy

## Abstract

Catheters are indispensable medical tools for accessing blood vessels, hollow organs, and body cavities to facilitate medication delivery and fluid drainage. However, they also serve as major entry points for bacterial contamination and trigger foreign body responses, necessitating locking strategies that are both bactericidal and biocompatible. This study introduces the first gel-based catheter lock, in contrast to conventional liquid locks. The gel is a poloxamer-based hydrogel formulated with 2-hydroxypropyl α-cyclodextrin (HP-αCD). HP-αCD forms supramolecular complexes with the poloxamer to enhance gelation, and with the nitric oxide (NO) donor to modulate NO release kinetics. This thixotropic gel can be injected into the catheter lumen when the catheter is not in use and withdrawn when vascular access is needed. The gel matrix provides a physical barrier that slows bacterial migration and minimizes drug loss. Simultaneously, the released NO functions as a broad-spectrum antimicrobial agent, effectively preventing biofilm formation on both the internal and external surfaces of the catheter. The NO-releasing hydrogel also demonstrates excellent hemocompatibility and reduces clot adhesion. Together, the gel-based lock offers a promising strategy for more effective catheter maintenance and represents a new application of hydrogels.

## 1 Introduction

Central venous catheters (CVCs) are an essential component of modern medical care, routinely used for hemodialysis, chemotherapy, and parenteral nutrition.^[1]^ While they provide immediate and reliable vascular access, CVCs are associated with two major complications: thrombus-related dysfunction and catheter-related bloodstream infections (CRBSIs).^[2]^ In the context of hemodialysis, catheter dysfunction occurs at a rate of approximately 0.5 to 3.4 episodes per 1,000 catheter-days, often necessitating catheter removal.^[3]^ According to the Centers for Disease Control and Prevention (CDC) Surveillance Summary of Bloodstream Infections in Outpatient Hemodialysis Facilities (2014–2019), 63% (98,502 out of 156,805) of reported bloodstream infections occurred in patients with CVCs.^[4]^ CRBSIs are a major contributor to hospitalization and mortality in this population and also impose a substantial financial burden, with estimated costs per episode ranging from $4,888 to $11,591.^[5]^

In response to the significant morbidity and healthcare burden associated with catheter occlusion and infection, catheter coating strategies have been developed to address these complications. Coatings such as pyrolytic carbon, albumin, elastin-like polypeptides, and heparin have shown promise in maintaining catheter patency, while antimicrobial coatings, such as chlorhexidine/silver sulfadiazine, minocycline/rifampin, and platinum/silver, have demonstrated efficacy in reducing infection risk.^[6,7]^ However, despite progress in both research and clinical translation, coating-based approaches face key limitations. First, a limited amount of drugs can be loaded into the thin coating. Second, the addition of surface coatings often leads to increased manufacturing costs, which may limit widespread clinical adoption. Third, coatings cannot be adjusted, replaced, or replenished during the course of treatment. Compared to catheter coatings, catheter lock technique provides superior cost efficiency and functional adaptability, enabling the use of formulations that can be modified or replaced as needed.^[8]^ When a CVC is not in use, its lumens are filled with lock solutions that are designed to maintain catheter patency.^[9]^ Anticoagulants such as heparin, ethylenediaminetetraacetic acid (EDTA), and trisodium citrate are commonly used in lock solutions to prevent and treat catheter thrombosis.^[6]^ In parallel, a wide range of antibiotics (e.g., vancomycin, gentamicin, ciprofloxacin) and antiseptics (e.g., alcohol, taurolidine, trisodium citrate) solutions have been employed for antimicrobial lock therapy.^[6,8,10]^ In clinical practice, it is common to combine anticoagulant and antimicrobial agents in a single lock formulation to address both major complications simultaneously. For example, in 2023, the U.S. Food and Drug Administration (FDA) approved DefenCath, a taurolidine-heparin (antiseptic + anticoagulant) lock solution, for the reduction of CRBSIs in adults with kidney failure undergoing chronic hemodialysis via CVCs.^[11]^

Although the catheter lock technique is a simple and practical approach to address both CVC-related complications, its broader application is limited by the unintended entry of lock solution into systemic circulation, even when the instilled volume does not exceed the priming volume of the catheter.^[12–15]^ Several studies have demonstrated that patients receiving heparin catheter locks after dialysis become systemically anticoagulated, with partial thromboplastin time values exceeding 200 seconds, far above the normal range of 25–35 seconds.^[12,16,17]^ Similarly, in a study involving ethanol lock therapy, 8 out of 9 patients experienced systemic adverse effects, including transient light-headedness, euphoria, and nausea, indicating ethanol leakage into the bloodstream.^[18]^ The leakage of lock solutions is primarily due to the absence of a physical barrier between the lock solution and contacting fluids. During instillation, a lock solution is injected to replace the pre-existing flush solution within the catheter lumen.^[19]^ Due to the flow characteristics of Newtonian fluids, approximately 15–20% of the lock solution is immediately spilled into the bloodstream when the injection volume equals the lumen capacity.^[14,15]^ Following instillation, the lock solution continues to be progressively lost over time due to the absence of any physical boundary between the lock and circulating blood.^[17,20]^ Therefore, it is conceptually intuitive that introducing a physical interface between a lock and its contacting fluid could mitigate leakage caused by unrestricted mixing.

Hydrogels have been widely utilized in biomedical and pharmaceutical fields, such as drug delivery and tissue engineering, due to their high water content, tunable mechanical properties, permeability, and excellent biocompatibility.^[21,22]^ With advances in biomaterials, a growing number of injectable hydrogels have emerged and are being explored in applications such as bioengineered 3D printing and localized drug administration.^[23]^ Given their semi-solid nature, injectable hydrogels may fulfill the aforementioned need by serving as an alternative to traditional liquid lock solutions, forming a boundary between a lock and other fluids to minimize catheter lock leakage. However, to the best of our knowledge, no hydrogels have been reported for use as catheter locks.

Herein, we present a Pluronic^®^ F127-based injectable and withdrawable hydrogel designed as a viable alternative to traditional catheter lock solutions. In light of the dual clinical needs of CVCs, we select S-nitrosoglutathione (GSNO)/2-hydroxypropyl α-cyclodextrin (HP-αCD) complex as a model therapeutic agent and incorporated it into the hydrogel matrix. Given the potent antimicrobial and antithrombotic properties of nitric oxide (NO),^[24–28]^ various NO-releasing compounds, including GSNO,^[29,30]^ low-molecular-weight N-diazeniumdiolates,^[31]^ S-nitroso-N-acetyl-penicillamine-conjugated ampicillin,^[32]^ and S-nitroso-N-acetyl-L-cysteine ethyl ester,^[33]^ have been previously added to lock solutions to mitigate the infectious and thrombotic complications of CVCs. However, all previous NO-releasing locks are based on solutions.

## 2 Results and Discussion

### 2.1 Formulation of NO-Releasing Hydrogels as Catheter Locks

A wide range of NO-releasing hydrogels have been developed using natural polymers such as chitosan, alginate, gelatin, and hyaluronic acid, as well as synthetic polymers including polyethylene glycol, polypropylene glycol, poly(acrylic acid), polyvinyl alcohol (PVA), and peptide amphiphiles.^[34–39]^ NO donors, including S-nitrosothiols, N-diazeniumdiolates, metal nitrosyl complexes, organic nitrates, and nitrite, are incorporated into these polymeric systems either through physical blending or chemical attachment to impart NO release capabilities. However, a majority of previous studies on NO-releasing hydrogels have focused on topical applications such as wound healing, treatment of skin infections, and enhancement of dermal blood flow. These applications impose markedly different requirements from those of catheter locks, particularly in terms of mechanical property, hemocompatibility, and NO release characteristics. For example, a key and unique requirement of a hydrogel-based catheter lock is that it must be easily injected into and withdrawn from a CVC without obstruction. F127 is an ideal candidate for this application due to its thermosensitivity and thixotropic properties.^[40,41]^ F127 is a triblock copolymer with a poly(ethylene oxide)−poly(propylene oxide)−poly(ethylene oxide) (PEO−PPO−PEO) structure. FDA has approved its use for oral, ophthalmic, and topical medicinal applications. At low temperatures, F127 remains in liquid state due to the high solubility of its PEO and PPO segments in water. As temperature increases, hydrophobic interactions among PPO chains become dominant, triggering self-assembly of copolymers into micelles. When the F127 concentration in the system is above the critical gelation concentration, these micelles organize into a structured network, resulting in a sol-to-gel transition. This phase transition is reversible. The micelle network disassembles upon cooling, allowing the hydrogel to return to its liquid state. The Oliveira group pioneered NO-releasing hydrogels using pure F127 as well as F127 mixed with polymers such as poly(acrylic acid) and PVA.^[42–44]^ F127-based hydrogels and films have been investigated for applications in wound healing and dermal vasodilation.^[42–48]^ Herein, we focus on the development of injectable and withdrawable F127 gels that are hemocompatible, minimize drug leakage, and release desirable levels of NO. Our gel formulations leverage supramolecular chemistry^[30]^ to achieve properties that make them well-suited as antimicrobial and antithrombotic locks for intravascular catheters.

Although F127 is well known to form thermoresponsive gels, the addition of a drug could perturb the gelation process. To determine the optimal NO-releasing F127 hydrogel formulation, the GSNO stock solution is blended with various F127 formulations in an ice bath, and a tube inversion test is performed after the solutions are incubated at room temperature (RT) and 37°C for 5 minutes. As shown in Figure 1A, in the presence of 0.1 M GSNO, at least 21.5 w/v% F127 is needed to form a hydrogel, even though the gelation of pure F127 only requires a concentration of 17 w/v%.^[49]^ Previously, we found that αCD and its derivatives can modulate NO release from GSNO solutions by forming GSNO/CD complexes.^[30]^ Herein, HP-αCD is added into the GSNO stock solution before they are mixed with the F127 solution. With an equimolar amount of HP-αCD and GSNO, the required F127 concentration for hydrogel formation at 37°C dropped to 20 w/v%. The CD-enhanced gelation occurs presumably because free αCD molecules (those not complexed with GSNO) can thread onto the flanking PEO segments of F127, forming PEO/αCD complexes in the solution state.^[50,51]^ As the temperature increases, the formation of microcrystalline PEO/αCD complexes ^[50,51]^ further enhances the sol-to-gel transition in addition to the original micelle-based gelation mechanism. Dynamic shear rheometer testing demonstrated the same effect of GSNO and CD (Figures 1B and S1). Although GSNO weakens the gel structure, as indicated by a reduction in storage modulus (G′, Figure 1B) and an increase in loss factor (tanδ, Table S1), the addition of an equimolar amount of HP-αCD compensated for this GSNO effect, recovering the gel integrity.

**Figure 1.**
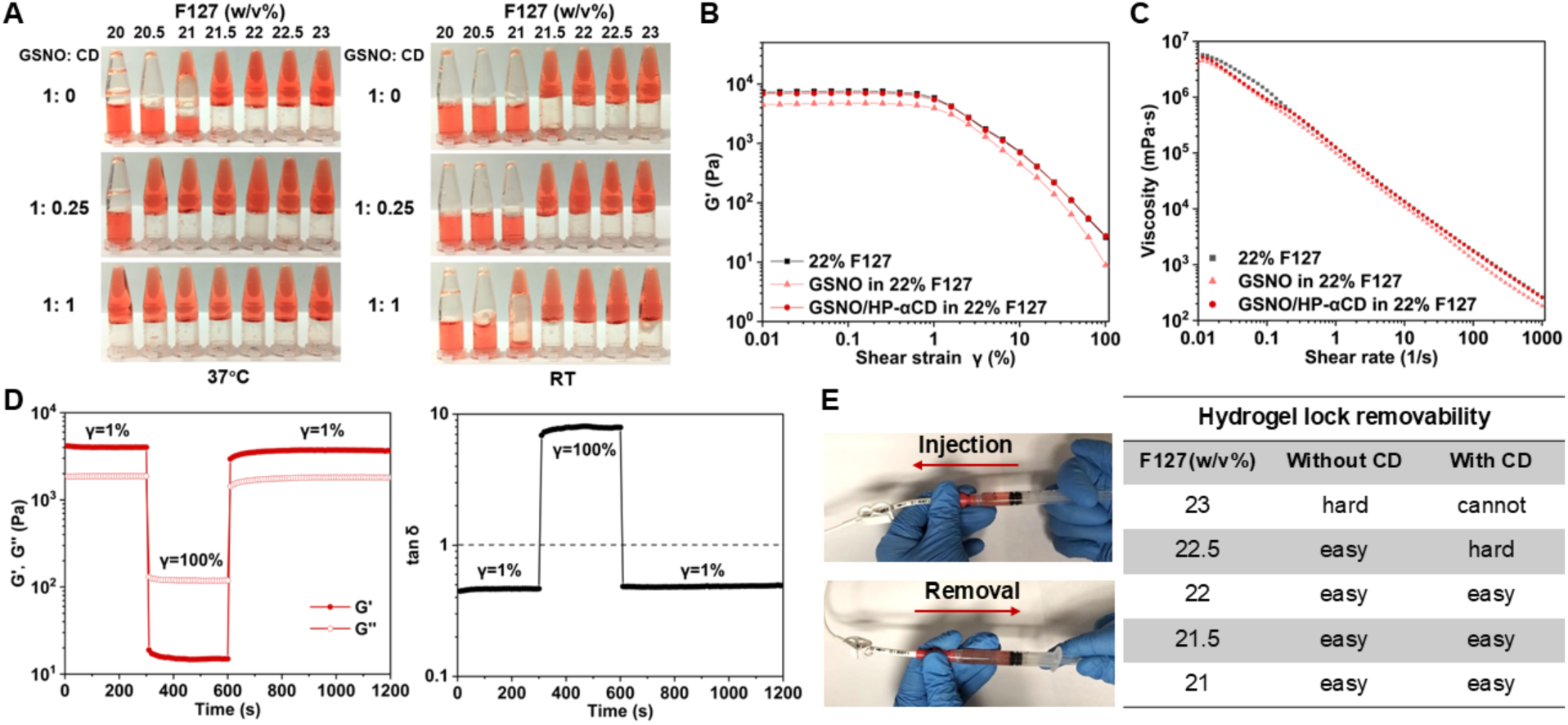
Physical characterization, rheological analysis, and functional screening of F127 hydrogels loaded with 0.1 M GSNO. (A) Tube inversion tests after incubating various F127 formulations at RT or 37°C for 5 min. (B) Storage modulus (G′) as a function of the strain amplitude at a frequency of 10 rad/s at 37 °C. (C) Shear thinning property at 37°C. (D) Thixotropic property of 22 w/v% F127 containing 0.1 M GSNO and 0.1 M HP-αCD at 37°C. (E) Injectability and removability of the gel evaluated in a central venous catheter.

The shear-thinning behavior of F127-based hydrogels arises from the reversible disruption of micellar structures under shear. This property allows the material to flow under applied force and is therefore essential to the injectability and removability of the catheter lock. As is shown in Figure 1C, incorporation of GSNO and HP-αCD did not compromise the overall shear-thinning property. Figure 1D shows the thixotropic behavior of the F127 gel loaded with GSNO and HP-αCD in a three-step shear test. The shear strain (γ) is successively switched each 300 s from a low amplitude of 1% to a high amplitude value of 100% and again to 1%. Upon application of a transient 100% shear strain, the gel exhibits liquid-like behavior, as indicated by an increase in tan δ to values greater than 1, and rapidly returns to its original state once the shear is removed (Figure 1D and S2). This shear-thinning and self-healing behavior suggest that the gel lock is both injectable and withdrawable under shear stress, while capable of reverting to a stable gel state after the cessation of mechanical force. We confirmed this injectability and removability using a commercial CVC with its indwelling portion placed in a 37°C water bath. All examined hydrogels can be easily infused into the catheter from a plastic syringe (Video 1). Hydrogels made of 22.5 w/v% F127 or less in the absence of HP-αCD and made of 22 w/v% F127 or less in the presence of HP-αCD allow obstruction-free aspiration (Figure 1E, Videos 2-4).

### 2.2 Spillage of Solutions and Gels as the Catheter Lock

To maintain patency and sterility of chronic CVCs, catheters are routinely filled with lock solutions containing heparin and, in some cases, antimicrobial agents.^[9]^ However, the clinical efficacy and safety of liquid-based catheter locks are frequently undermined by drug leakage into the bloodstream.^[13,15]^ This spillage partially arises due to the laminar flow profile of Newtonian fluids within the catheter, where fluid at the center moves faster than at the periphery (Figure 2A, left). Moreover, before instilling a drug-loaded lock, the catheter lumen has already been filled with a flush solution, which is usually saline.^[19]^ When a lock solution is instilled to displace the saline flush, the absence of physical barrier between the lock solution and the saline leads to immediate and inevitable mixing of liquids and then drug loss to the bloodstream. Experimental and theoretical models indicate that up to 25% of the lock solution can enter circulation upon instilling a volume equal to the catheter’s internal volume, with spillage beginning as early as 50% of lumen filling.^[14,15,17,52]^ The speed of manual injection has minimal to no effect on the extent of instillation spillage.^[52]^ Unlike low-viscosity aqueous solutions, the hydrogel exhibits piston-like behavior during injection due to the higher viscosity even under the shear stress and is also less prone to mixing with saline. As a result, chemical spillage out of the catheter is significantly reduced when a gel instead of a solution is being injected as a lock (Figure 2A). Beyond instillation, additional loss can occur via gravitational sinking and concentration gradient–driven diffusion, particularly for dense or drug-rich lock formulations.^[17,20]^ Clinical evidence corroborates such inevitable drug loss with reported systemic adverse effects, including heparin-induced thrombocytopenia, citrate-induced hypocalcemia, ethanol-related neurological symptoms, and gentamicin-associated ototoxicity.^[12,16,18,53]^ This fundamental limitation of the liquid lock is expected to be mitigated by switching to a hydrogel-based lock. The hydrogel does not easily sink into the blood and the gel network constrains drug molecules, thereby reducing gradual loss of the drug into the bloodstream after instillation.

**Figure 2.**
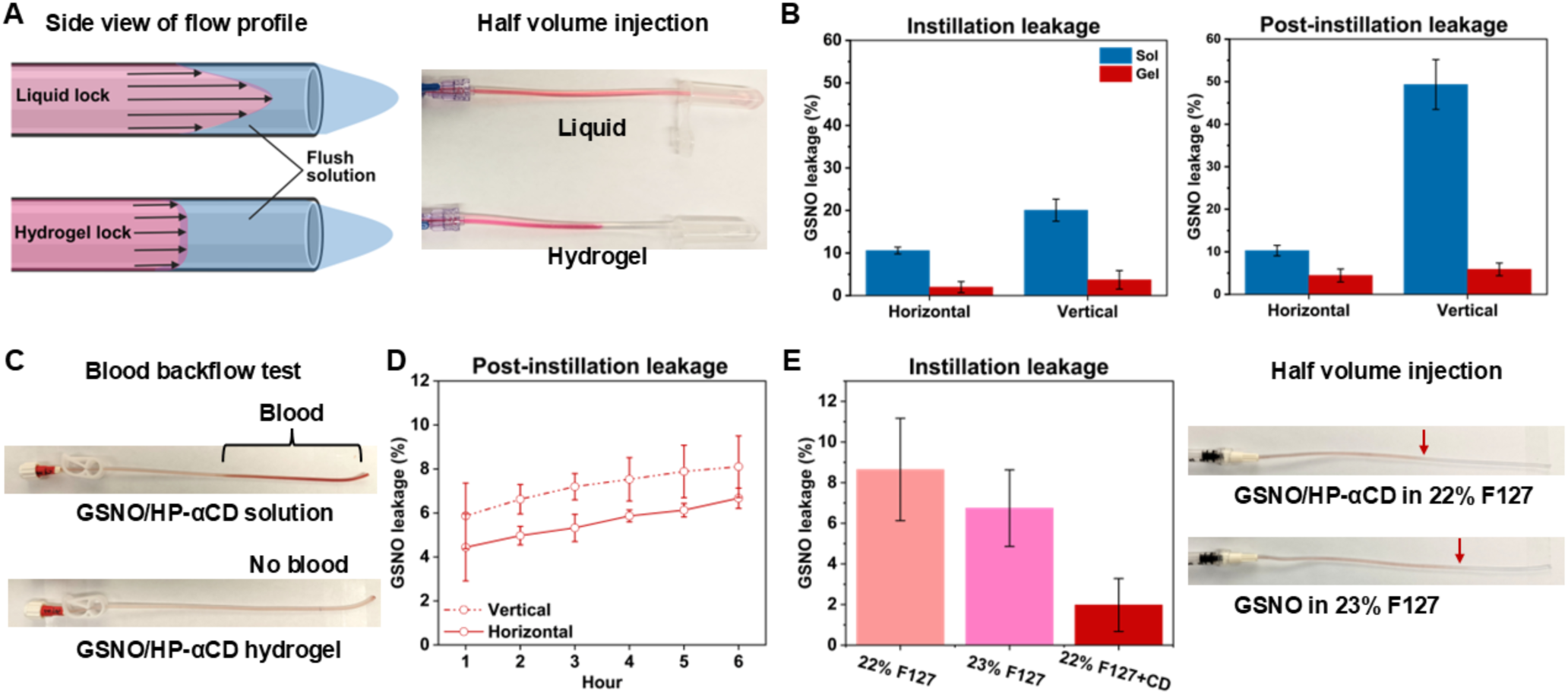
*In vitro* evaluation of the leakage of the liquid- and gel-based catheter locks. (A) Schematic illustration and a photo showing the flow behavior of liquid and gel locks injected into catheters prefilled with a flush solution (saline). The lock contains a red dye to aid visualization. (B) Quantification of GSNO leakage during instillation (n=5) and 1 hour after instillation (n=3) for liquid lock versus 22 w/v% F127 hydrogel lock under horizontal and vertical catheter orientations. All formulations contain 0.1 M GSNO and 0.1 M HP-αCD. (C) Photos showing the blood backflow toward the liquid lock instead of the hydrogel lock. (D) GSNO leakage from the hydrogel lock over 6 h under horizontal and vertical catheter orientations (n=3). (E) GSNO leakage during horizontal instillation of different lock formulations (0.1 M GSNO in 22 w/v% F127 gel, 0.1 M GSNO in 23 w/v% F127 gel, and 0.1 M GSNO/0.1 M HP-αCD in 22 w/v% F127 gel). The red arrow indicates the approximate interface between the lock and the pre-filled saline.

To evaluate this hypothesis, we conducted *in vitro* tests to comprehensively evaluate the leakage of solutions and gels as the catheter locks, incorporating both horizontal and vertical orientations to simulate different clinical scenarios based on patient positioning. The National Kidney Foundation Kidney Disease Outcomes Quality Initiative clinical practice guidelines recommends that CVC tips should be placed in the mid-to-deep right atrium to reduce the risk of catheter malfunction.^[54]^ When the CVC tip is positioned in the right atrium and surrounded by flowing blood, the gravitational influence on the lock varies depending on whether the patient is in a reclined or upright position during lock instillation. Subsequent patient movements when the catheter is not in use also influence lock leakage over time. The catheter orientation significantly affects the extent of leakage for the solution-based lock formulation. When pushing the prefilled saline with the GSNO/HP-αCD lock solution at a volume equal to that of the catheter lumen, horizontal infusion results in 10.6±0.8% leakage, whereas vertical infusion doubles the leakage (Figure 2B, left). In contrast, only less than 4% of GSNO is spilled during the instillation of the hydrogel regardless of the orientation (Figure 2B, right). When a catheter filled with the GSNO/HP-αCD solution is suspended vertically with the tip immersed in PBS, nearly 50% of the loaded drug leaked within just 1 hour (Figure 2B, right). Positioning the catheter horizontally reduced such post-instillation leakage to 10.3±1.2% (Figure 2B, right). Similarly, blood backflow is observed when the catheter tip is vertically immersed in blood due to the exchange of the GSNO/HP-αCD solution with blood (Figure 2C). This phenomenon is primarily attributed to the high density of the GSNO/HP-αCD solution (∼1.166 g/mL). In contrast to the liquid lock, the GSNO/HP-αCD-loaded F127 hydrogel as the lock effectively mitigates these issues, as no blood backflow is observed under the same experimental conditions (Figure 2C) and no more than 6% GSNO leaks one hour past the instillation (Figure 2B). Even after 6 hours, only less than 8% of GSNO diffuses away from the gel lock (Figure 2D), indicating that the hydrogel network effectively retains the loaded drug. Notably, the GSNO-loaded 23 w/v% F127 hydrogel without HP-αCD suffers from significantly more leakage than that in 22 w/v% F127 with HP-αCD (Figure 2E), even though their viscoelastic properties are similar (Figures S1 and S2). Pure F127 hydrogel is known for its relatively high tendency to dissolve in aqueous solutions.^[55]^ The formation of supramolecular HP-αCD/PEO domains presumably reduces the gel dissolution in the pre-filled saline during the instillation, thereby minimizing spillage. Based on these findings, 22 w/v% F127 hydrogel containing 0.1 M GSNO and 0.1M HP-αCD was selected for all subsequent studies.

To further prove the reduced spillage of the gel-based catheter lock relative to conventional solution-based lock, we leverage the potent vasodilatory properties of NO as a physiological indicator of intravascular leakage.^[56]^ The buffer and F127 hydrogel containing GSNO/HP-αCD are alternately instilled into the same rat, with each formulation administered three times. As shown in Figure 3, each instillation of the solution elicits a rapid and significant decrease in the mean arterial pressure (MAP), indicating immediate systemic exposure to NO due to the GSNO spillage. In contrast, no such hemodynamic response is observed following administration of the hydrogel-based formulation, suggesting minimal leakage into circulation.

**Figure 3.**
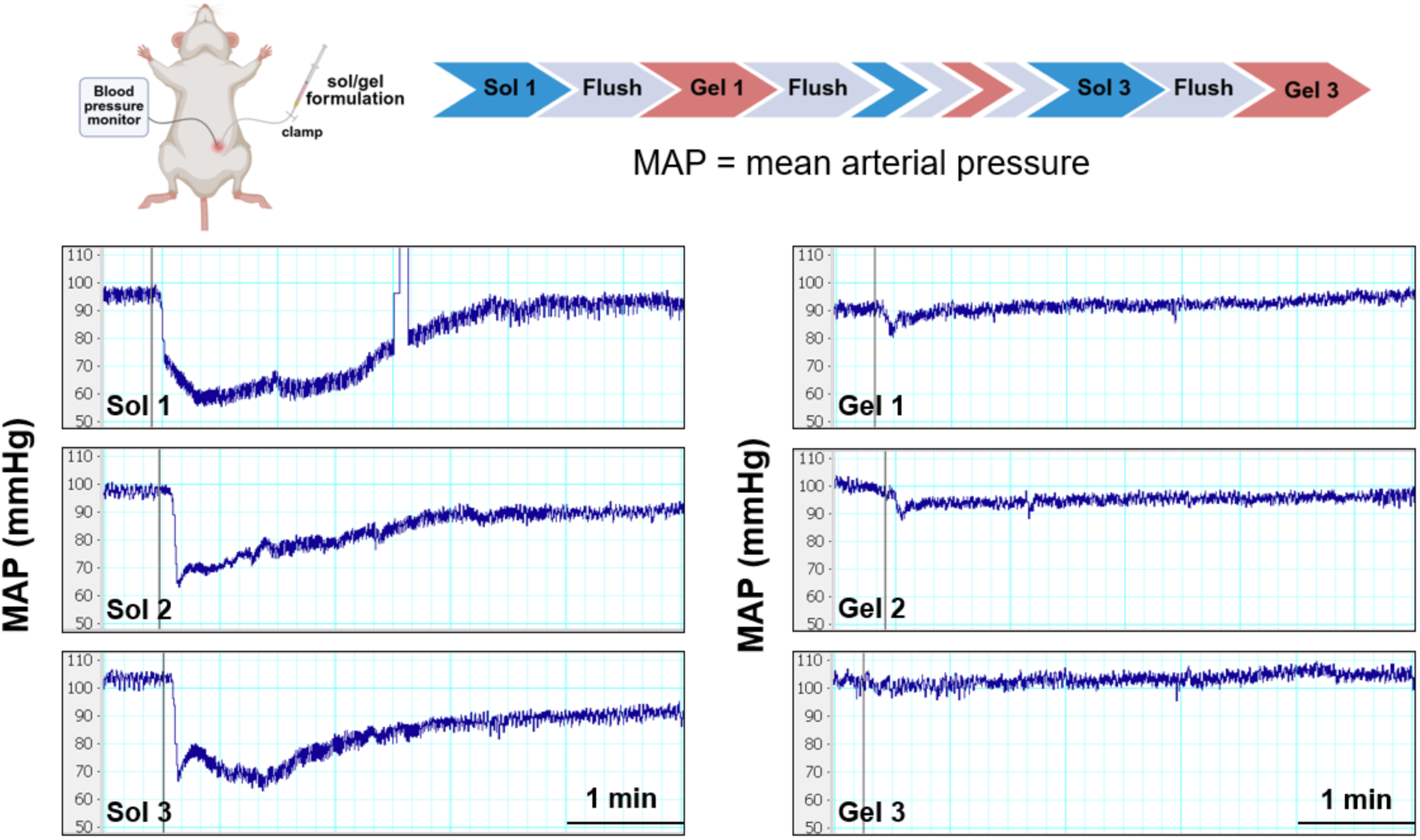
*In vivo* assessment of GSNO spillage from the solution-based or hydrogel-based catheter lock using a rat model. The blood pressure drop is caused by NO released from the spilled GSNO during instillation. The vertical gray lines indicate when the lock is injected.

### 2.3 Hemocompatibility and cytotoxicity of the NO-Releasing Gel Lock

Considering that the lock at the CVC tip is in direct contact with blood, hemocompatibility of the lock medium is a critical safety requirement: the catheter lock should not induce clot formation at the gel-blood interface; the hydrogel should be dissolvable if accidentally introduced into the bloodstream; the lock should not cause significant hemolysis. In our initial efforts to devise hydrogel-based catheter locks, we screened multiple hydrogel candidates such as those based on synthetic silicate nanoplatelets and PVA. While silicate nanoplatelet-based hydrogel demonstrated excellent shear-thinning behavior, it shows a strong proclivity to induce blood clot formation at the hydrogel–blood interface. *In vitro* clotting assays show that blood in contact with 6 wt% silicate nanoplatelet hydrogel (Figure 4A, PC) clots within 3 minutes, significantly faster than the control group (Figure 4A, C, blood only), which clots after 6 minutes. This procoagulant effect is attributed to electrostatic interactions between the charged nanoplatelet surfaces and the blood components.^[57]^ Similar clotting behavior has been observed with other clay-based materials, such as kaolin. Due to this procoagulant risk, this group of inorganic hydrogels were excluded from further consideration as the catheter lock. In contrast, the F127 hydrogel does not induce clotting earlier than the control (Figure 4A, F127). More interestingly, no clot adhesion was observed on the surface of the NO-releasing F127 hydrogel over the course of the 6-min assay (Figure 4A, NO). This aligns with the well-known anti-coagulant role of NO because it inhibits platelet aggregation and adhesion.^[58–62]^ Although this study does not focus on comprehensive evaluation of the anticoagulant function of NO, the NO-releasing hydrogel is expected to exhibit both antimicrobial and antithrombotic properties to mitigate two major complications of CVC, namely infection and thrombosis. This dual-acting property is a highly unique advantage of NO over other drugs used in catheter locks.

**Figure 4.**
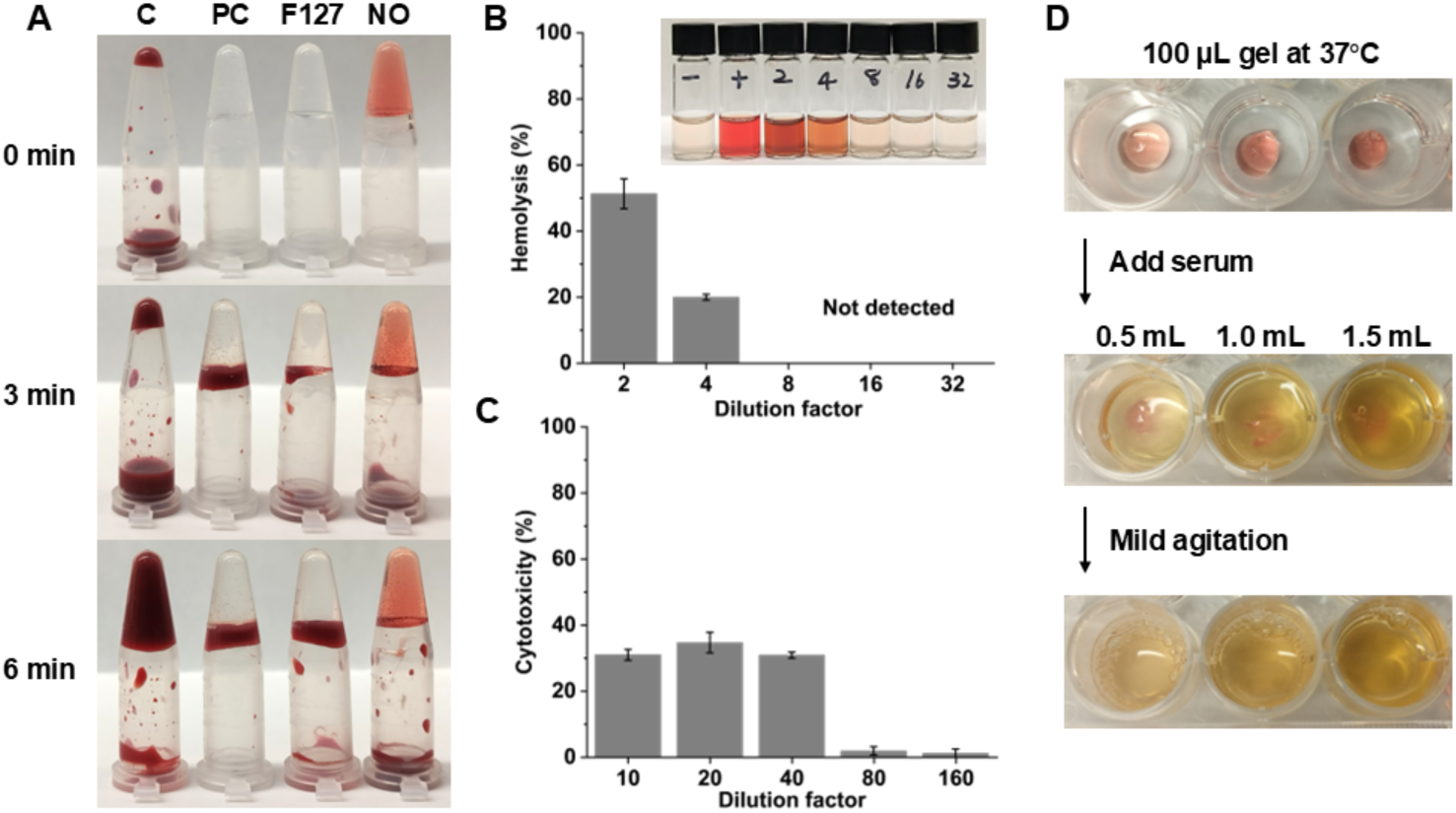
Hemocompatibility of the NO-releasing hydrogel lock. (A) Blood clotting tests at 37 °C. C: control (blood only); PC: positive control (6 wt% silicate nanoplatelet gel); F127: 22 w/v% F127 gel; NO: 0.1 M GSNO/0.1 M HP-αCD in 22 w/v% F127 gel. (B) Hemolysis tests of 22 w/v% F127 gel containing 0.1 M GSNO and 0.1 M HP-αCD. (C) Cytotoxicity tests of the same hydrogel. (D) Photos showing dissolution of the GSNO-HP-αCD-loaded F127 hydrogel in serum.

Hemolysis assays reveal that the GSNO/HP-αCD hydrogel does not lyse red blood cells when diluted 8-fold or more (Figure 4B). The hydrogel shows no detectable cytotoxicity when diluted 80-fold or more (Figure 4C). Given the small volume of the catheter lumen (up to 3 mL) relative to the total human blood volume (∼5 liters) and the limited leakage of the catheter lock, the hydrogel is unlikely to pose hemolytic or cytotoxic risk. Another important consideration is whether the hydrogel lock can dissolve in blood in case it inadvertently enters the circulation, as insoluble materials may pose a risk of vascular blockage. During our material screening process, hydrogels based on PVA were excluded because even small debris of these hydrogels cannot get dissolved in serum at physiological temperature. In contrast, the F127 hydrogel can be dissolved in serum within 1 minute of mild physical agitation (Figure 4D). Given the continuous flow of blood, the F127 hydrogel formulation will not cause any blood vessel blockage.

### 2.4 NO Release of the Hydrogel-Based Catheter Locks

GSNO is a naturally occurring NO donor that spontaneously decomposes at body temperature to release NO and generate glutathione disulfide (GSSG).^[56]^ To investigate the NO release kinetics, the decomposition of GSNO with or without HP-αCD in the F127 hydrogel at 37°C is monitored based on the characteristic UV-Vis absorption of GSNO (Figure 5A). In the absence of HP-αCD, approximately 73% of GSNO decomposes within the first 24 hours. In agreement with our previous observations in solution-based systems,^[30]^ the addition of HP-αCD slows down the GSNO degradation due to the formation of host-guest complexes. When an equimolar amount of HP-αCD is added, the GSNO concentration after 24 hours of incubation at 37 °C is nearly doubled compared to the hydrogel without HP-αCD. Furthermore, the lifetime of GSNO is extended from 3 to 5 days due to the presence of HP-αCD.

**Figure 5.**
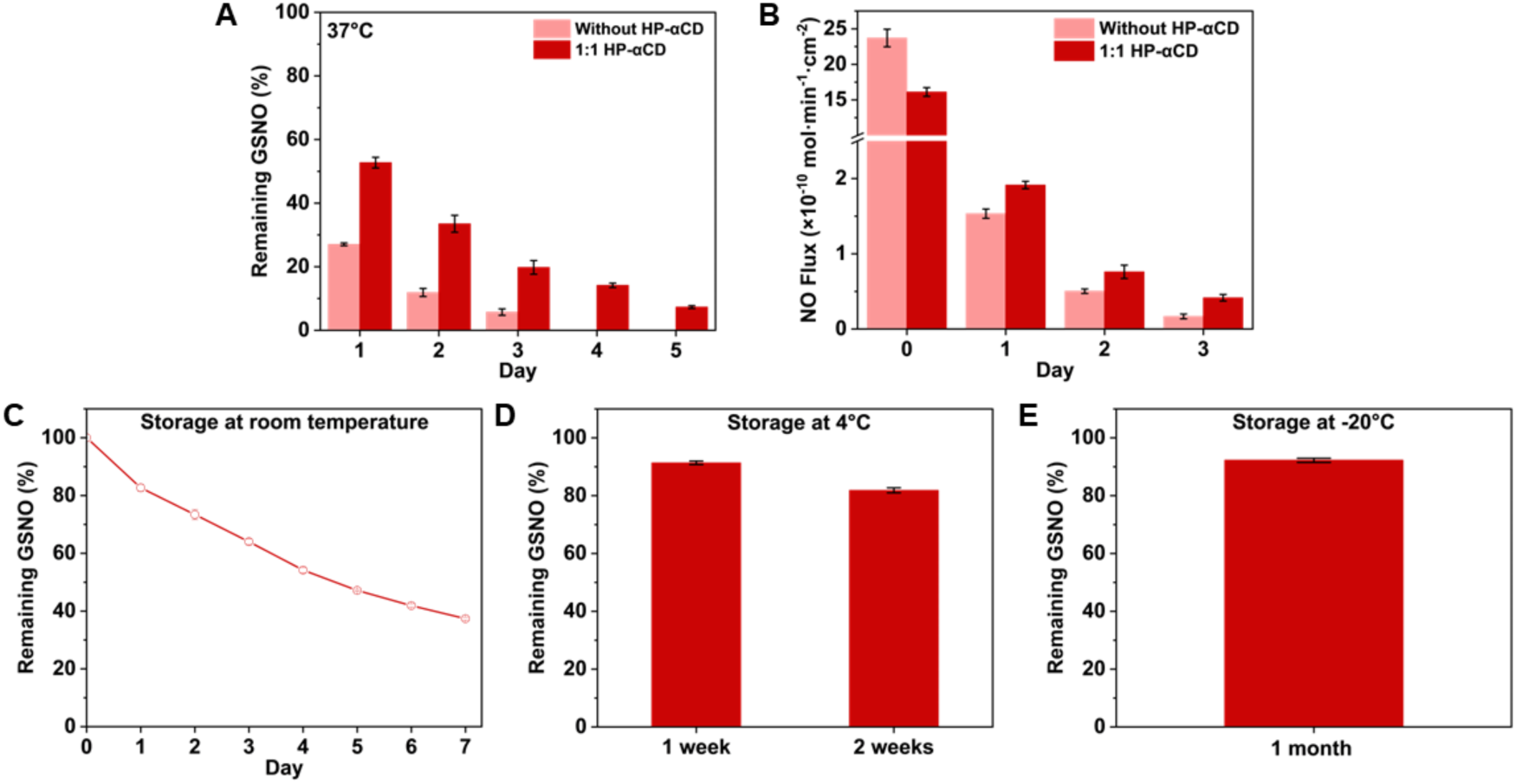
Stability and NO release property of the GSNO-loaded F127 hydrogel. (A) Decomposition of GSNO at 37 °C in 22 w/v% F127 hydrogel with or without equimolar HP-αCD. (B) NO flux measured from the outer surface of sealed catheters filled with GSNO-loaded F127 gels with and without HP-αCD. (C–E) Storage stability of the 22 w/v% F127 hydrogel containing 0.1 M GSNO and 0.1 M HP-αCD at RT, 4 °C, and −20 °C. All experiments were performed in triplicate.

Another distinct advantage of using NO as the antimicrobial agent is that NO can permeate through the polymeric wall of the catheter and thus protect both the intraluminal and extraluminal environments. Figure 5B shows the NO release from sealed catheter segments filled with GSNO-loaded hydrogel formulations with and without HP-αCD. In the absence of HP-αCD, the initial NO flux exceeds 23 × 10⁻¹⁰ mol min⁻¹ cm⁻². Inclusion of HP-αCD suppresses this undesirable initial burst release to approximately 16 × 10⁻¹⁰ min⁻¹ mol cm⁻². During the subsequent tests over three days, the NO flux from the HP-αCD-containing hydrogel remains consistently higher than that of the CD-free hydrogel. For CVCs used for hemodialysis, catheter locks are typically refreshed every 48 to 72 hours.^[63]^ Therefore, the NO release profile of the hydrogel lock aligns well with these lock replacement intervals.

### 2.5 Storage Stability of the Hydrogel Lock

To evaluate the storage stability of the F127 hydrogel containing 0.1 M GSNO and 0.1 M HP-αCD, samples are stored at RT, 4°C, and -20°C in the dark. After 24 hours at RT, 17.3% of GSNO in hydrogel has decomposed. The GSNO half-life in this hydrogel formulation is about 5 days at RT (Figure 5C). When stored at 4°C, the characteristic absorbance of GSNO remains unchanged over 24 hours. Approximately 92% and 82% of GSNO is left after one week and two weeks of storage, respectively (Figure 5D). Freezing the hydrogel further improves the storage stability. After one month of storage at -20°C, only 7.8% of GSNO decomposes (Figure 5E). Additionally, the hydrogel retains its thermoresponsive gelation properties after storage in the refrigerator or freezer (data not shown). These findings suggest that the hydrogel lock may be stored at low temperatures.

### 2.6 Antibacterial Properties of the NO-Releasing Catheter Lock

#### 2.6.1 Hydrogel-based catheter lock reduces microbial migration

The prevention and treatment of CRBSIs remain a significant clinical challenge for patients undergoing hemodialysis via a CVC. Approximately 70% of dialysis-related bloodstream infections occur in patients using catheters.^[4]^ *Staphylococcus* species are the leading pathogens that cause CRBSIs, with *Staphylococcus* aureus (*S.* aureus) accounting for 21–43% of cases, and methicillin-resistant *S.* aureus (MRSA) reported in 12–38% of cases.^[64,65]^ The primary source of catheter infection is closely linked to the duration of catheter use. In short-term cases (<10 days), infections typically originate from cutaneous organisms colonizing the external surface of the catheter, whereas in the long-term use (>10 days), infection is more often due to intraluminal spread from the catheter hub (Figure 6A).^[66,67]^

**Figure 6.**
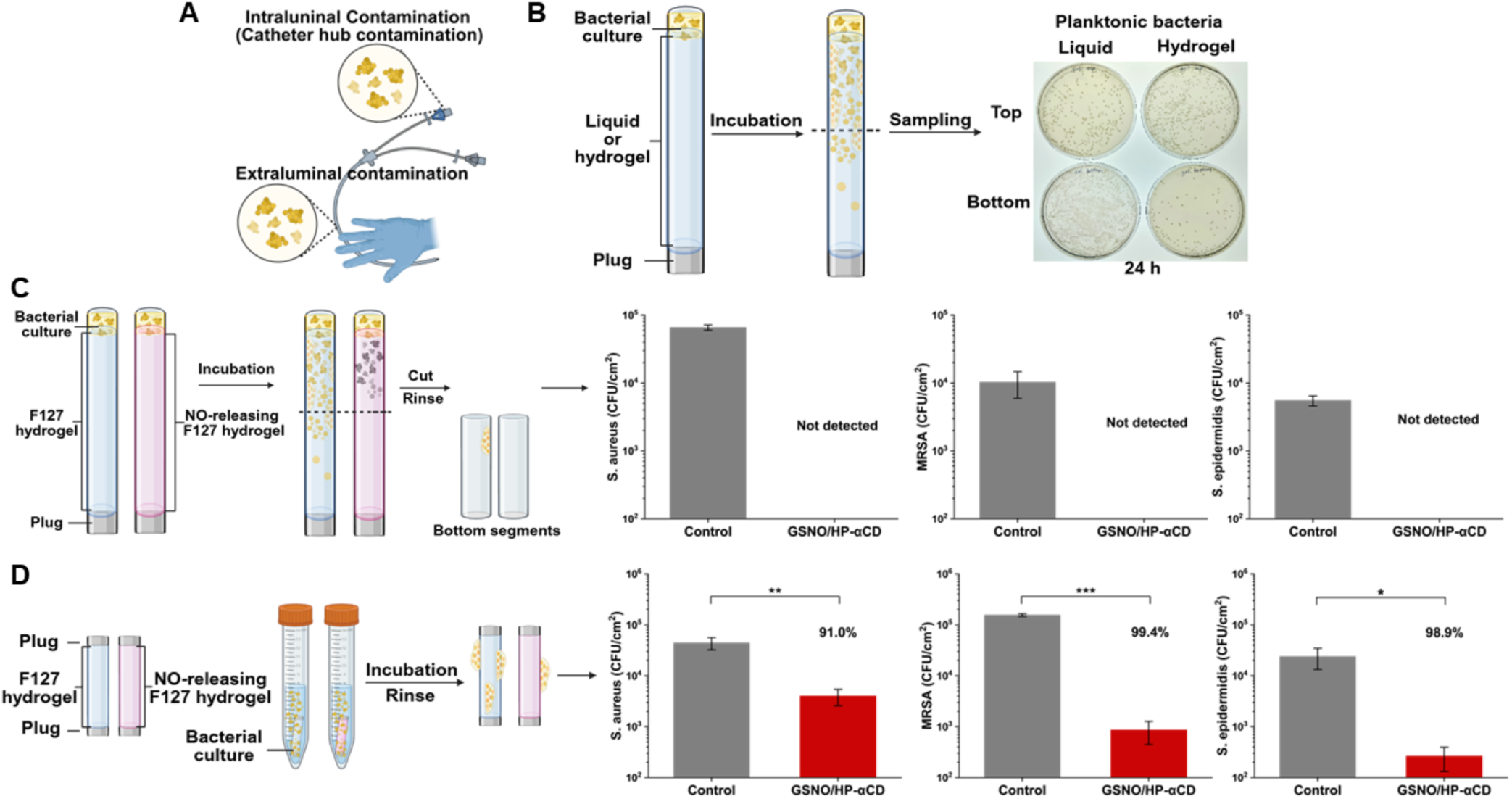
NO-releasing gel locks reduce bacterial growth. (A) Schematic illustration of potential routes of bacterial contamination on intravascular catheters. (B) Comparison of *S. aureus* migration in catheters filled with a liquid or hydrogel medium. (C) 3-day biofilm tests on the intraluminal surface of the catheter filled with a NO-releasing F127 hydrogel and a control hydrogel. and (D) 3-day biofilm tests on the extraluminal surface of the catheter. The percent reduction is indicated on each graph. All experiments were performed in triplicate. *p < 0.05; **p < 0.01; **p < 0.001.

While the liquid lock allows bacteria to move freely, the hydrogel lock provides a much more rigid physical matrix that slows down bacterial migration from the catheter hub to the distal end of the catheter. An *in vitro* bacterial migration model illustrated in Figure 6B was used to evaluate the effectiveness of the F127 hydrogel in impeding bacterial movement along the catheter. 10 µL of *S. aureus* suspension is gently added to the opening of a catheter tube filled with a solution or a hydrogel to mimic hub contamination. After 1-day incubation at 37°C, the planktonic bacteria in the top half and the bottom half of the catheter lock are quantified. As shown in Figure 6B, for liquid-filled catheters, *S. aureus* accumulates in the bottom segment because of facile diffusion and sedimentation. In contrast, the planktonic bacteria are much less in the bottom part of the catheter lock when the lock is a gel, indicating that the hydrogel-based lock as a rigid medium reduces bacterial mobility. However, F127 hydrogel alone cannot completely block bacterial migration over time without being combined with antimicrobial agents. The continuous growth of bacterial biofilm along the inner surface of the catheter will ultimately lead to infection. We confirmed that significant bacterial biofilms are formed throughout the catheter over the period of 3 days (Figure S3), necessitating the use of a antibacterial drug in the catheter lock.

#### 2.6.2 NO-releasing gel lock prevents intraluminal biofilm formation

For patients with long-term catheter use and a history of CRBSIs, clinical practice guidelines recommend the use of highly concentrated antibiotic lock solutions, typically 100 to 1,000 times higher than the minimal inhibitory concentration (MIC).^[10]^ However, as discussed above, leakage of such high-concentration antibiotics into the bloodstream is inevitable, raising concerns about systemic side effects and the promotion of antibiotic resistance. NO is a highly reactive free radical with potent antimicrobial properties.^[68–71]^ It quickly reacts with oxygen and superoxide from the microbiological environment to produce reactive oxygen and nitrogen species, which place oxidative and nitrosative stress on microbes.^[24,25]^ Unlike traditional antibiotics that typically act through a single mechanism, NO exhibits multifaceted antimicrobial activities, ranging from enzyme deactivation and lipid peroxidation, to membrane disruption and direct damage to microbial DNA and DNA repair systems.^[24,25,72]^ These broad and overlapping mechanisms make NO a potent, broad-spectrum antimicrobial agent. We compared the viable bacterial biofilm attached to the bottom catheter segment when the catheter is filled with the F127 hydrogel with and without GSNO/HP-αCD. Three *Staphylococcus* strains, commonly implicated in CRBSIs, were tested as representative pathogens. After three days of incubation, although the drug-free hydrogel lock allows for significant growth of bacteria, no viable bacteria can be detected from the catheter segments filled with the NO-releasing hydrogel, across all three strains (Figure 6C). The substantial reduction in biofilm formation demonstrates that the NO-releasing hydrogel lock is highly effective in preventing intraluminal bacterial colonization.

#### 2.6.3 NO-releasing gel lock reduces extraluminal biofilm formation

While the majority of CRBSIs originate from hub contamination, a significant portion of CRBSIs are associated with bacterial colonization on the extraluminal surface.^[66,67]^ These extraluminal infections typically result from improper handling during catheter implant or from patient skin flora entering the body through the insertion site.^[66]^ Therefore, preventing bacterial colonization along the outer surface of the catheter is also important for comprehensive infection control. Traditional antibacterial agents in lock solutions are confined within the catheter lumen due to their negligible diffusivity in the catheter wall. Therefore, they cannot target bacteria colonizing the external surface of the catheter. In contrast, NO is a tiny gaseous molecule capable of diffusing through polymeric catheter materials, offering a unique advantage in addressing extraluminal infections. This non-direct-contact antimicrobial protection provided by NO-releasing locks has been previously reported by the Brisbois Group, the Schoenfisch group, and our group.^[30,31,33]^ To evaluate the effectiveness of the NO-releasing gel lock in preventing extraluminal infections, catheter segments filled with hydrogel are sealed at both ends and incubated with the three representative *Staphylococcus* strains. On the third day, the biofilm formed on the outer surface of the catheter is quantified. As shown in Figure 6D, 1- to 2-log reductions in biofilm are obtained on NO-releasing catheters compared to the control catheters. Specifically, *S. aureus*, MRSA, and *S. epidermidis* exhibit a 91.0%, 99.4%, and 98.9% reduction, respectively, confirming antibacterial protection on the extraluminal catheter surface provided by the NO generated from the hydrogel and diffused across the polymeric catheter wall.

#### 2.6.4 NO-releasing hydrogel treats catheter infections

If a CVC is already contaminated, antibiotic lock therapy may be employed to eradicate established biofilms that form on the internal surface of the catheter.^[12,73]^ However, antibiotics often possess limited effectiveness against established biofilms, largely due to the extracellular polymeric substances (EPS) that impede drug penetration into the biofilm.^[72]^ In contrast, NO is capable of diffusing through the EPS matrix, disrupting the biofilm structure and exposing the bacteria.^[72]^ We evaluated the biofilm eradication efficacy of the NO-releasing hydrogel lock. Catheter segments are sealed at one end, inoculated with three types of bacterial cultures, respectively, and incubated for 24 hours to allow biofilm formation on the intraluminal surface. After incubation, the bacterial culture medium is removed, and the catheter lumen is filled with the hydrogel with or without GSNO/HP-αCD, followed by an additional 24-hour incubation. After the 1-day treatment, reduced biofilm growth is observed across all three tested strains (Figure 7A). The *S. epidermidis* biofilm shows the most dramatic response to NO, with viable cell counts reduced by ∼2 orders of magnitude. The biofilm biomass of *S. aureus* and MRSA decreases to 16.5% and 5% of the untreated controls, respectively.

**Figure 7.**
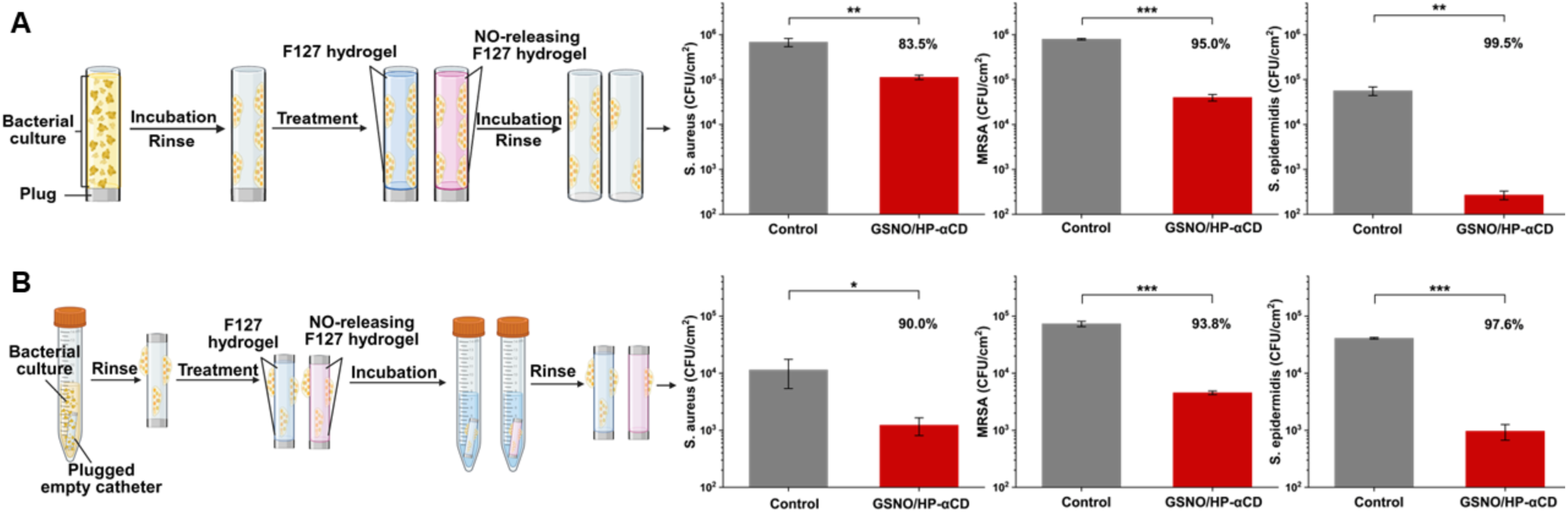
NO-releasing hydrogel lock eradicates established bacterial biofilms on inner (A) and outer (B) catheter surfaces. Established biofilms are treated with the F127 hydrogel lock containing 0.1 M GSNO and 0.1 M HP-αCD for 24 hours. The percent reduction is indicated on each graph. All experiments were performed in triplicate.*p < 0.05; **p < 0.01; **p < 0.001.

As discussed previously, current antibiotic lock therapies are ineffective against infections occurring on the outer surface of catheters because organic molecules cannot readily diffuse across the catheter wall. To assess whether our NO-releasing hydrogel lock could disperse and kill bacteria on the outer surface of the catheter, the hydrogel is added into the catheter segment that has established bacterial biofilms on the external surface. Both ends of the catheter segment are sealed so NO can only reach the outer surface via the catheter wall. Remarkably, despite the biofilms being located on the opposite side of the catheter wall, the NO-releasing hydrogel is still able to substantially reduce mature biofilm (Figure 7B). Following the 24-hour treatment, there is at least 90% reduction in biofilm biomass compared to controls for all three bacterial strains. These results underscore the highly unique efficacy of NO as a diffusive small-molecule drug in preventing and treating catheter-associated infections.

## 3 Conclusions

Catheter locks based on liquid formulations often spill into the bloodstream during instillation due to the parabolic flow pattern and subsequently mix with blood owing to the absence of a defined interface. When the lock solution contains drugs such as antimicrobial and anticoagulant agents, this spillage can lead to systemic toxicity and reduced therapeutic efficacy. In contrast, the use of a hemocompatible thixotropic hydrogel as a catheter lock significantly minimizes spillage and mixing, thereby slowing drug loss. When combined with GSNO, the hydrogel effectively inhibits bacterial growth and migration through sustained NO release. Future work may explore alternative hydrogel systems to fine-tune thixotropic properties, viscosity, stability, and NO release kinetics. Additional functions of NO-releasing hydrogel locks, such as inhibition of platelet aggregation and modulation of inflammatory responses, will also be systematically examined in future studies. Beyond central venous catheters, this hydrogel lock concept may also be extended to other types of catheters, such as peritoneal dialysis catheters and biliary drainage catheters.

## 4 Materials and Methods

### 4.1 Chemicals and reagents

Poloxamer 407 (Pluronic^®^ F127) was purchased from Spectrum Chemical Mfg. Corp. Sodium phosphate dibasic (Na_2_HPO_4_), sodium hydroxide (NaOH), L-glutathione reduced (GSH), sodium nitrite, fetal bovine serum (FBS), and penicillin/streptomycin were purchased from MilliporeSigma. Luria−Bertani (LB) broth powder and agar were purchased from Thermo Fisher Scientific. HP-αCD was purchased from Cyclodextrin-Shop. Laponite-XLG is a gift from BYK USA lnc. Tryptic soy broth (TSB) was purchased from BD biosciences. Eagle’s Minimum Essential Medium (EMEM), murine fibroblast L929 and bacterial strains including S. aureus (25923), methicillin-resistant S. aureus (MRSA, BAA-2312), and S. epidermidis (12228) were purchased from the American Type Culture Collection (ATCC).

### 4.2 GSNO synthesis and characterization

GSNO was synthesized by nitrosating GSH in an acidic nitrite solution. In brief, 4.59 grams (14.94 mmoles) of GSH were dissolved in 29.87 mL of 0.5 M hydrochloric acid (14.94 mmoles). The mixture was stirred at 0°C for 10 minutes. Subsequently, 1.03 grams (14.94 mmoles) of sodium nitrite was added, and the reaction was stirred at the same temperature for 40 minutes, ensuring the flask was protected from light. Afterward, 10 mL of cold acetone was added to the mixture and stirred for an additional 10 minutes. The GSNO precipitate was collected through vacuum filtration and thoroughly washed with cold deionized water. Finally, the GSNO was freeze-dried and stored in the dark at -20°C until further use. GSNO was characterized by ^1^H NMR (400 MHz, DMSO) and ^13^C NMR (100 MHz, DMSO) spectroscopy to confirm its structure; full spectra and peak assignments are provided in the supplementary Information (Figure S4).

### 4.3 Preparation of NO-releasing solutions and hydrogels

HP-αCD powder was weighed and dissolved in 0.1 M Na₂HPO₄ to make a 0.5 M HP-αCD stock solution. Then, 168.16 mg of GSNO was added to 1 mL of the HP-αCD stock, and NaOH was gradually introduced to adjust the pH to 7.4 and facilitate dissolution of GSNO. The Pluronic F127 stock solution (29 w/v%) was prepared using a standard cold method. Briefly, F127 powder was slowly added to 0.1 M cold phosphate buffer (pH = 7.4) under continuous stirring at 4 °C, and the mixture was stirred overnight until a clear, homogeneous solution was obtained. For the solution-based formulations, the GSNO/HP-αCD stock was diluted 5-fold with the phosphate buffer. For the hydrogel formulations, the GSNO/HP-αCD stock was blended with 29 w/v% F127 hydrogel in its liquid state and diluted with phosphate buffer in an ice bath to obtain the 0.1 M concentration of GSNO and HP-αCD as well as different final concentrations of F127.

### 4.4 Rheological measurements

The rheological tests were conducted using an Anton Paar MCR 702e rheometer equipped with a Peltier device for precise temperature control. A parallel plate geometry was used, featuring a 25 mm diameter upper plate and a fixed gap of 0.5 mm. A solvent trap was used throughout the experiments to prevent solvent loss by evaporation and preserve sample hydration. The shear viscosity (η) of the gels was measured under steady shear flow conditions across a range of shear rates (γ͘) from 0.01 to 1000 s⁻¹. Amplitude sweep tests were conducted on gels maintained at 37 °C to identify the upper strain limit (γL) of the linear viscoelastic region and to determine the yield stress (σ₀). Self-healing tests were performed using strain step oscillatory measurements at a fixed angular frequency (ω) of 10 rad/s. The strain amplitude was alternated every 300 seconds, starting with a small strain (1%) in the linear viscoelastic region, followed by a large strain (100%) within the nonlinear regime, and then returned to 1% to assess recovery. The elastic modulus (G′) and viscous modulus (G″) were recorded throughout to evaluate the energy stored and dissipated during each deformation cycle and to monitor structural recovery.

### 4.5 Lock leakage evaluations

To evaluate lock leakage under conditions mimicking clinical use, an *in vitro* catheter model was developed to mimic the essential features of a standard CVC, including a proximal connector compatible with a Luer-lock syringe, a flexible catheter body, a functional clamp, and an end cap.

#### 4.5.1 Leakage during instillation

The catheter was pre-filled with bubble-free saline and clamped to mimic a clinical catheter flush procedure. A luer-lock syringe was filled with a solution or hydrogel formulation at a volume matching the internal volume of the catheter. The syringe with the hydrogel was warmed in a 37 °C incubator for 5 minutes to enable the gelation. The Luer-lock syringe was then connected to the connector of the catheter. The catheter was positioned either horizontally or vertically. After opening the clamp, the formulation was slowly and steadily infused into the catheter. The fluid flowing out of the distal end of the catheter was collected in a 1.5 mL microcentrifuge tube for subsequent quantification.

The percentage of GSNO leakage is defined as the ratio of the GSNO detected from the microcentrifuge tube to that infused into the catheter. The total infused GSNO was calculated from the known formulation concentration (0.1 M) and the injection volume of 0.22 mL. The leaked GSNO was quantified by measuring the absorbance of the collected fluid at 335 nm in a 96-well plate using a Varioskan™ LUX Multimode Microplate Reader. By multiplying the measured concentration by the collected volume in the microcentrifuge tube, the amount of leaked GSNO was obtained. Each formulation was tested with n = 5 replicates.

#### 4.5.2 Leakage post instillation

The catheter was first locked with a solution or hydrogel. The clamp was closed and the proximal end of the catheter was capped. The catheter was then placed in a 37 °C incubator, either horizontally or vertically. The catheter tip was immersed in 1 mL of PBS solution in a tube that was sealed to prevent evaporation. At designated time points, the PBS was sampled to quantify the spilled GSNO via absorbance at 335 nm. The GSNO leakage was calculated as described above. Each formulation was tested with n = 3 replicates.

#### 4.5.3 Blood backflow assay

The blood backflow assay followed the same procedure as described for the post-instillation leakage test, with the catheter tip immersed in 1 mL of rat blood containing 4% (w/v) sodium citrate instead of PBS. The catheters were hanged vertically in the incubator.

### 4.6 *In vivo* rat experiment to estimate the lock leakage

An adult male Sprague Dawley rat (318 g, Envigo) was anesthetized via inhalation of isoflurane (1–2% in medical-grade oxygen) under spontaneous breathing. The anesthetic depth was adjusted to the minimal level necessary to abolish spinal and canthal reflexes. A polyethylene catheter (PE-50 tubing, ADInstruments) was inserted into the femoral artery for continuous monitoring of hemodynamic variables, including MAP, recorded using a PowerLab data acquisition system (ADInstruments). Playback of the MAP data were used after the experiment for data analysis of blood pressure responses. A second PE catheter (Scientific Commodities, Inc., #BB31695-PE/5) was inserted into the femoral vein for testing of experimental catheter lock solution and gel formulations. After hemodynamic baseline values were observed, 0.1 mL of GSNO/HP-αCD-loaded lock solutions or gels were alternately placed into the venous catheter using a 1-ml syringe. After each lock solution/gel placement, the blood pressure response was recorded until a new stable baseline was reached. The catheter was flushed with a total of 0.5 mL of 5 units/mL heparin in LR (Lactated Ringer’s solution) before another solution or gel was tested. The rat was humanely euthanized with an anesthetic overdose at the end. The experiment was approved by the Virginia Commonwealth University (VCU) Institutional Animal Care and Use Committee (AD10003237).

### 4.7 Hemocompatibility and cytotoxicity tests

#### 4.7.1 Blood clotting tests

Four 1.5 mL microcentrifuge tubes were placed in a 37 °C dry bath. One tube was left empty and the other three tubes were each filled with 0.4 mL of hydrogels: 6 w/v% silicate nanoplatelet hydrogel, 22 w/v% F127 hydrogel, and GSNO/HP-αCD-loaded 22 w/v% F127 hydrogel. To restore coagulation activity in sodium citrate (4 w/v%) -treated rat blood, 0.02 mL of 0.2 M CaCl₂ was added to each mL of blood. Subsequently, 0.6 mL of calcium-supplemented blood was added to the empty tube, and 0.2 mL was added to each of the hydrogel-containing tubes. The clot formation was monitored by inverting the tubes at 1-minute intervals.

#### 4.7.2 Hydrogel solubility in serum

0.1 mL of GSNO/HP-αCD-loaded 22 w/v% F127 hydrogel was transferred into the wells of a 24-well plate preheated on a 37 °C hot plate. Pre-warmed serum at volumes of 0.5, 1.0, and 1.5 mL was then added to individual wells containing the hydrogel. The plate was gently agitated by hand, and the dissolution behavior of the hydrogel was visually monitored.

#### 4.7.3 Hemolysis test

Rat blood was centrifuged at 1,000 × g for 10 minutes to separate the red blood cells (RBCs). The supernatant was discarded, and the RBC pellet was resuspended in PBS to prepare a 10% (v/v) RBC suspension. The NO-releasing hydrogel was mixed with PBS and RBC suspension to reach 2-, 4-, 8-, 16-, and 32 -fold dilutions. The final RBC concentration is 5 % in these microcentrifuge tubes. PBS and deionized water were used as negative and positive controls, respectively. After 1h of incubation at 37 °C, the sample was centrifuged at 1,000 × g for 5 minutes, and the absorbance (540 nm) of the collected supernatant was measured by a microplate reader (Varioskan™ LUX Multimode Microplate Reader). Since GSNO has a pink color that absorbs at 540 nm, the final absorbance value was corrected by subtracting the background absorbance of GSNO at the corresponding concentration in PBS. Hemolysis (%) was calculated using the following formula:

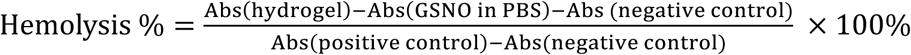

#### 4.7.4 Cytotoxicity

L929 murine fibroblast cells were cultured in complete media (EMEM with 1% penicillin/streptomycin and 10% FBS) before being seeded in a 96-well plate at a concentration of 10^4^ cells per well for 24 h. Then, the culture medium was aspirated, and cells were exposed to the 10-, 20-, 40-, 80-, and 160-fold diluted NO-releasing hydrogel or EMEM. After 24 h incubation, cytotoxicity was assessed using a lactate dehydrogenase (LDH) leakage assay following the manufacturer’s protocol (CyQUANT^TM^ LDH Cytotoxicity Assay C20301). Unexposed cells served as a negative control, and cells exposed to lysis buffer served as a positive control. Absorbance at 490 nm (reference 680 nm) was measured by Varioskan™ LUX Multimode Microplate Reader, and cytotoxicity was calculated relative to positive and negative controls.

### 4.8 Quantification of GSNO decomposition via UV–Vis absorption spectroscopy

GSNO/HP-αCD F127-based hydrogels were placed in 1.5 mL disposable polystyrene cuvettes with caps and stored at different temperatures in the absence of light. Absorbance at 545 nm was measured using a UV-Vis spectrophotometer (Go Direct® Fluorescence/UV-VIS Spectrophotometer). If air bubbles were observed, the cuvettes were placed on ice to temporarily convert the hydrogel to its liquid state, allowing trapped air to be released before measurements. All measurements were performed in triplicate.

### 4.9 Measurement of NO release from the catheter locks

A chemiluminescence NO analyzer (ECO PHYSICS nCLD 66) was used to monitor NO diffusion from the outer surface of catheters. Medical-grade silicone catheters (HelixMark® 60-011-07; 1.58 mm ID, 2.41 mm OD) were cut into 2.5 cm segments, filled with NO-releasing liquid or hydrogel formulations, and tightly sealed at both ends using plastic rod plugs. The sealed catheter segments were immersed in 4 mL of PBS at 37 °C in an amber glass sample cell. The NO analyzer was calibrated using N₂ and NO gas of a known concentration. During measurements, N₂ gas was continuously purged into the sample cell at a flow rate of 100 cm³/min to carry the released NO into the chemiluminescence detector. Between measurements, all samples were stored separately in 5 mL of PBS at 37 °C in the dark.

### 4.10 Antibacterial tests

All three *Staphylococcus* species used in this study were maintained on LB agar plates and stored at 4 °C. For each experiment, fresh colonies cultured within 48 hours were used to ensure viability and consistency. The bacterial inoculum was standardized using a 0.5 McFarland turbidity standard (approximately 1.5 × 10⁸ CFU/mL), and all working suspensions were prepared by diluting the 0.5 McFarland suspension in the appropriate culture medium by a factor of 10,000. All medical-grade silicone catheters (HelixMark® 60-011-07), plastic rods, F127 stock solution, phosphate buffer, and other experimental apparatus were sterilized by autoclaving prior to use.

#### 4.10.1 Intraluminal bacterial migration tests

Catheters were cut into 4.5 cm long segments. The bottom end of the tube was tightly sealed with a plastic rod. A solution or 22 w/v% F127 hydrogel containing 1% TSB is added into the tube, leaving ∼10 µL of space at the open end. 10 µL of 10^4^-fold diluted McFarland bacterial suspension in 1% TSB was added to the open end to mimic hub contamination. Each tube was then placed vertically into a sterile culture tube containing sterile PBS at a level just below the open end to maintain a humid condition. After 24h-incubation at 37 °C, 10 μL of liquid or gel was collected from the top and bottom segments of the tube, respectively, for planktonic bacteria quantification via plate counting. The sample from the top segment was taken after discarding the uppermost 10 μL medium. For biofilm quantification, each tube with the F127 hydrogel was cut in half. The hydrogel was carefully aspirated, and the catheter was gently rinsed with sterile PBS to remove loosely associated bacteria. Then, each segment of tube was transferred into a centrifuge tube containing 2 mL of sterile PBS and subjected to vigorous vortexing for 1 minute to dislodge biofilm bacteria. 20 µL of the resulting bacterial suspension was plated on LB agar plates for plate counting. The final results were expressed as colony-forming units per square centimeter of inner surface area of the tube (CFU/cm²).

#### 4.10.2 Intraluminal biofilm prevention study

The protocol is similar to the biofilm quantification protocol detailed in 4.10.1. The catheter was filled with F127 hydrogel with or without GSNO/HP-αCD. After bacterial inoculation, all tubes were incubated at 37 °C for 3 days before biofilm quantification for bottom segments.

#### 4.10.3 Extraluminal biofilm prevention study

Catheter tubes (2.5 cm) were filled with F127 hydrogel or NO-releasing F127 hydrogel and sealed at both ends by plastic rods. The sealed catheters were immersed in 2 mL of 1% TSB containing the 10^4^-diluted 0.5 McFarland bacterial suspension. The bacterial culture medium is refreshed every 24 hours. After 72 hours, the tubes were retrieved for biofilm quantification on the external catheter surface.

#### 4.10.4 Biofilm eradication study

To evaluate the biofilm eradication capability of the NO-releasing hydrogel, biofilm of Staphylococcus strains was first established on the catheter. For the intraluminal surface, 2.5 cm tubes were sealed at one end and filled with a bacterial suspension in TSB (∼10^4^ CFU/mL). For the extraluminal surface, 2.5 cm catheter segments sealed at both ends were fully immersed in 2 mL of the same bacterial suspension. All samples were incubated at 37 °C for 24 hours to allow for formation of mature biofilms on the internal or external surface. For the intraluminal biofilm eradication study, the bacterial culture medium was removed from the tube and the lumen was filled with F127 hydrogel with or without GSNO/HP-αCD. For the extraluminal biofilm eradication study, one end of the tube was unplugged to allow addition of the F127 hydrogel with or without GSNO/HP-αCD. Then the tubes were transferred into sterile PBS. All tubes were further incubated at 37 °C for additional 24 hours before tube segments were processed for biofilm quantification.

#### 4.10.5 Statistical analysis

All antibacterial tests were triplicated. Data were presented as mean ± SD. Statistical analysis was performed using the Student’s t-test. A p-value <0.05 was considered statistically significant.

## Acknowledgements

We acknowledge financial support from the NIH (R01HL168133) and the Virginia Commonwealth University (CCTR Endowment Fund).

## Competing interests

The authors declare no competing interests.

## Data availability

Additional data that support the findings of this study are available from the corresponding author upon reasonable request.

## Notes

### Competing Interest Statement

The authors have declared no competing interest.

## References

(1) Kehagias, E.; Galanakis, N.; Tsetis, D. Central Venous Catheters: Which, When and How. British Journal of Radiology 2023, 96 (1151). 10.1259/bjr.20220894.

(2) Wang, Y.; Sun, X. Reevaluation of Lock Solutions for Central Venous Catheters in Hemodialysis: A Narrative Review. Renal Failure 2022, 44 (1), 1501–1518. 10.1080/0886022X.2022.2118068.

(3) Beathard, G. Malfunction of chronic hemodialysis catheters - UpToDate. 2025. https://www.uptodate.com/contents/malfunction-of-chronic-hemodialysis-catheters

(4) CDC. Surveillance Summary of Bloodstream Infections in Outpatient Hemodialysis Facilities — National Healthcare Safety Network, 2014-2019. 2014. https://www.cdc.gov/dialysis-safety/media/pdfs/BSI-NHSN-2014to2019-508.pdf

(5) Harron, K.; Mok, Q.; Hughes, D.; Muller-Pebody, B.; Parslow, R.; Ramnarayan, P.; Gilbert, R. Generalisability and Cost-Impact of Antibiotic-Impregnated Central Venous Catheters for Reducing Risk of Bloodstream Infection in Paediatric Intensive Care Units in England. PLoS One 2016, 11 (3). 10.1371/JOURNAL.PONE.0151348,.

(6) CDC. Background Information: Strategies for Prevention of Catheter-Related Infections in Adult and Pediatric Patients. 2011. https://www.cdc.gov/infection-control/hcp/intravascular-catheter-related-infection/prevention-strategies.

(7) Sousa, C.; Henriques, M.; Oliveira, R. Mini-Review: Antimicrobial Central Venous Catheters – Recent Advances and Strategies. Biofouling 2011, 27 (6), 609–620. 10.1080/08927014.2011.593261.

(8) Kim, E. Y.; Saunders, P.; Yousefzadeh, N. Usefulness of Anti-Infective Lock Solutions for Catheter-Related Bloodstream Infections. Mount Sinai Journal of Medicine 2010, 77 (5), 549–558. 10.1002/MSJ.20213.

(9) Niyyar, V. D. Catheter Dysfunction: The Role of Lock Solutions. Seminars in Dialysis 2012, 25 (6), 693–699. 10.1111/j.1525-139X.2011.00991.x

(10) Justo, J. A.; Bookstaver, P. B. Antibiotic Lock Therapy: Review of Technique and Logistical Challenges. Infection and Drug Resistance 2014, 7, 343–363. 10.2147/IDR.S51388

(11) FDA. FDA approves new drug under special pathway for patients receiving hemodialysis. 2023. https://www.fda.gov/drugs/news-events-human-drugs/fda-approves-new-drug-under-special-pathway-patients-receiving-hemodialysis

(12) Labriola, L.; Pochet, J. M. Any Use for Alternative Lock Solutions in the Prevention of Catheter-Related Blood Stream Infections? The Journal of Vascular Access 2017, 18, S34–S38. 10.5301/JVA.5000681.

(13) Mokrzycki, M. H.; Lok, C. E. Traditional and Non-Traditional Strategies to Optimize Catheter Function: Go with More Flow. Kidney International 2010, 78 (12), 1218–1231. 10.1038/KI.2010.332.

(14) Ash, S. R. Fluid Mechanics and Clinical Success of Central Venous Catheters for Dialysis-Answers to Simple but Persisting Problems. Seminars in Dialysis 2007, 20 (3), 237–256. 10.1111/j.1525-139X.2007.00284.x.

(15) Polaschegg, H. D.; Shah, C. Overspill of Catheter Locking Solution: Safety and Efficacy Aspects. ASAIO Journal 2003, 49 (6), 713–715. 10.1097/01.MAT.0000094040.54794.2D,.

(16) Karaaslan, H.; Peyronnet, P.; Benevent, D.; Lagarde, C.; Rince, M.; Leroux-Robert, C. Risk of Heparin Lock-Related Bleeding When Using Indwelling Venous Catheter in Haemodialysis. Nephrology Dialysis Transplantation 2001, 16 (10), 2072–2074. 10.1093/NDT/16.10.2072.

(17) Markota, I.; Markota, D.; Tomic, M. Measuring of the Heparin Leakage into the Circulation from Central Venous Catheters - An in Vivo Study. Nephrology Dialysis Transplantation 2009, 24 (5), 1550–1553. 10.1093/NDT/GFN696.

(18) Maiefski, M.; Rupp, M. E.; Hermsen, E. D. Ethanol Lock Technique: Review of the Literature. Infection Control & Hospital Epidemiology 2009, 30 (11), 1096–1108. 10.1086/606162.

(19) Goossens, G. A. Flushing and Locking of Venous Catheters: Available Evidence and Evidence Deficit. Nurse Research Practice 2015, 2015 (1), 985686. 10.1155/2015/985686.

(20) Polaschegg, H. D. Loss of Catheter Locking Solution Caused by Fluid Density. ASAIO Journal 2005, 51 (3), 230–235. 10.1097/01.MAT.0000159742.15560.93.

(21) Jacob, S.; Nair, A. B.; Shah, J.; Sreeharsha, N.; Gupta, S.; Shinu, P. Emerging Role of Hydrogels in Drug Delivery Systems, Tissue Engineering and Wound Management. Pharmaceutics 2021, 13 (3), 357. 10.3390/PHARMACEUTICS13030357.

(22) Lei, L.; Bai, Y.; Qin, X.; Liu, J.; Huang, W.; Lv, Q. Current Understanding of Hydrogel for Drug Release and Tissue Engineering. Gels 2022, 8 (5), 301. 10.3390/GELS8050301.

(23) Bertsch, P.; Diba, M.; Mooney, D. J.; Leeuwenburgh, S. C. G. Self-Healing Injectable Hydrogels for Tissue Regeneration. Chemical Reviews 2023, 123 (2), 834–873. 10.1021/acs.chemrev.2c00179.

(24) DeGroote, M. A.; Fang, F. C. Antimicrobial Properties of Nitric Oxide. Nitric Oxide and Infection 2002, 231–261. 10.1007/0-306-46816-6_12.

(25) De Groote, M. A.; Fang, F. C. NO Inhibitions: Antimicrobial Properties of Nitric Oxide. Clinical Infectious Diseases 1995, 21. 10.1093/CLINIDS/21.SUPPLEMENT_2.S162.

(26) Mendelsohn, M. E.; O’Neill, S.; George, D.; Loscalzo, J. Inhibition of Fibrinogen Binding to Human Platelets by S-Nitroso-N-Acetylcysteine. Journal of Biological Chemistry 1990, 265 (31), 19028–19034. 10.1016/S0021-9258(17)30619-1.

(27) Wo, Y.; Brisbois, E. J.; Bartlett, R. H.; Meyerhoff, M. E. Recent Advances in Thromboresistant and Antimicrobial Polymers for Biomedical Applications: Just Say Yes to Nitric Oxide (NO). Biomaterials Science 2016, 4 (8), 1161–1183. 10.1039/C6BM00271D.

(28) Wu, M.; Lu, Z.; Wu, K.; Nam, C.; Zhang, L.; Guo, J. Recent Advances in the Development of Nitric Oxide-Releasing Biomaterials and Their Application Potentials in Chronic Wound Healing. Journal of Materials Chemistry B 2021, 9 (35), 7063–7075. 10.1039/D1TB00847A.

(29) Ashcraft, M.; Douglass, M.; Garren, M.; Mondal, A.; Bright, L. E.; Wu, Y.; Handa, H. Nitric Oxide-Releasing Lock Solution for the Prevention of Catheter-Related Infection and Thrombosis. ACS Applied Bio Materials 2022, 5 (4), 1519–1527. 10.1021/acsabm.1c01272.

(30) Li, W.; Wang, D.; Lao, K. U.; Wang, X. Inclusion Complexation of S-Nitrosoglutathione for Sustained Nitric Oxide Release from Catheter Surfaces: A Strategy to Prevent and Treat Device-Associated Infections. ACS Biomaterials Science Engineering 2023, 9 (3), 1694–1705. 10.1021/acsbiomaterials.2c01284.

(31) Maloney, S. E.; Grayton, Q. E.; Wai, C.; Uriyanghai, U.; Sidhu, J.; Roy-Chaudhury, P.; Schoenfisch, M. H. Nitric Oxide-Releasing Hemodialysis Catheter Lock Solutions. ACS Applied Materials Interfaces 2023, 15 (24), 28907–28921. 10.1021/acsami.3c02506.

(32) Chug, M. K.; Griffin, L.; Garren, M.; Tharp, E.; Nguyen, G. H.; Handa, H.; Brisbois, E. J. Antimicrobial Efficacy of a Nitric Oxide-Releasing Ampicillin Conjugate Catheter Lock Solution on Clinically-Isolated Antibiotic-Resistant Bacteria. Biomaterial Science 2023, 11 (19), 6561–6572. 10.1039/D3BM00775H.

(33) Kumar, R.; Massoumi, H.; Chug, M. K.; Brisbois, E. J. S-Nitroso-N-Acetyl-l -Cysteine Ethyl Ester (SNACET) Catheter Lock Solution to Reduce Catheter-Associated Infections. ACS Applied Materials Interfaces 2021, 13 (22), 25813–25824. 10.1021/acsami.1c06427.

(34) Estes Bright, L. M.; Wu, Y.; Brisbois, E. J.; Handa, H. Advances in Nitric Oxide-Releasing Hydrogels for Biomedical Applications. Current Opinion in Colloid Interface Science 2023, 66, 101704. 10.1016/J.COCIS.2023.101704.

(35) Tavares, G.; Alves, P.; Simões, P. Recent Advances in Hydrogel-Mediated Nitric Oxide Delivery Systems Targeted for Wound Healing Applications. Pharmaceutics 2022, 14 (7). 10.3390/PHARMACEUTICS14071377.

(36) Yang, T.; Zelikin, A. N.; Chandrawati, R.; Yang, T.; Chandrawati, R.; Zelikin, A. N. Progress and Promise of Nitric Oxide-Releasing Platforms. Advanced Science 2018, 5 (6), 1701043. 10.1002/ADVS.201701043.

(37) Zheng, G.; Li, R.; Wu, P.; Zhang, L.; Qin, Y.; Wan, S.; Pei, J.; Yu, P.; Fu, K.; Meyerhoff, M. E.; Liu, Y.; Zhou, Y. Controllable Release of Nitric Oxide from an Injectable Alginate Hydrogel. International Journal of Biological Macromolecules 2023, 252, 126371. 10.1016/J.IJBIOMAC.2023.126371.

(38) Feura, E. S.; Maloney, S. E.; Conlon, I. L.; Broberg, C. A.; Yang, F.; Schoenfisch, M. H. Injectable Polysaccharide Hydrogels as Localized Nitric Oxide Delivery Formulations. Advanced Materials Technologies 2023, 8 (7), 2201529. 10.1002/admt.202201529.

(39) Younis, M.; Tabish, T. A.; Firdharini, C.; Aslam, M.; Khair, M.; Anjum, D. H.; Yan, X.; Abbas, M. Self-Assembled Peptide-Based Fibrous Hydrogel as a Biological Catalytic Scaffold for Nitric Oxide Generation and Encapsulation. ACS Applied Materials Interfaces 2025, 17 (19), 27964–27973. 10.1021/acsami.5c03250.

(40) Shamma, R. N.; Sayed, R. H.; Madry, H.; El Sayed, N. S.; Cucchiarini, M. Triblock Copolymer Bioinks in Hydrogel Three-Dimensional Printing for Regenerative Medicine: A Focus on Pluronic F127. Tissue Engineering Part B Reviews 2022, 28 (2), 451–463. 10.1089/ten.TEB.2021.0026.

(41) Li, S.; Yang, C.; Li, J.; Zhang, C.; Zhu, L.; Song, Y.; Guo, Y.; Wang, R.; Gan, D.; Shi, J.; Ma, P.; Gao, F.; Su, H. Progress in Pluronic F127 Derivatives for Application in Wound Healing and Repair. International Journal of Nanomedicine 2023, 18, 4485–4505. 10.2147/IJN.S418534.

(42) Champeau, M.; Póvoa, V.; Militão, L.; Cabrini, F. M.; Picheth, G. F.; Meneau, F.; Jara, C. P.; de Araujo, E. P.; de Oliveira, M. G. Supramolecular Poly(Acrylic Acid)/F127 Hydrogel with Hydration-Controlled Nitric Oxide Release for Enhancing Wound Healing. Acta Biomaterialia 2018, 74, 312–325. 10.1016/J.ACTBIO.2018.05.025.

(43) Vercelino, R.; Cunha, T. M.; Ferreira, E. S.; Cunha, F. Q.; Ferreira, S. H.; De Oliveira, M. G. Skin Vasodilation and Analgesic Effect of a Topical Nitric Oxide-Releasing Hydrogel. Journal of Materials Science: Materials in Medicine 2013, 24 (9), 2157–2169. 10.1007/s10856-013-4973-7.

(44) Schanuel, F. S.; Raggio Santos, K. S.; Monte-Alto-Costa, A.; de Oliveira, M. G. Combined Nitric Oxide-Releasing Poly(Vinyl Alcohol) Film/F127 Hydrogel for Accelerating Wound Healing. Colloids and Surfaces B Biointerfaces 2015, 130, 182–191. 10.1016/J.COLSURFB.2015.04.007.

(45) Cao, J.; Su, M.; Hasan, N.; Lee, J.; Kwak, D.; Kim, D. Y.; Kim, K.; Lee, E. H.; Jung, J. H.; Yoo, J. W. Nitric Oxide-Releasing Thermoresponsive Pluronic F127/Alginate Hydrogel for Enhanced Antibacterial Activity and Accelerated Healing of Infected Wounds. Pharmaceutics 2020, 12 (10), 926. 10.3390/PHARMACEUTICS12100926.

(46) Pelegrino, M. T.; de Araújo, D. R.; Seabra, A. B. S-Nitrosoglutathione-Containing Chitosan Nanoparticles Dispersed in Pluronic F-127 Hydrogel: Potential Uses in Topical Applications. Journal of Drug Delivery Science and Technology 2018, 43, 211–220. 10.1016/J.JDDST.2017.10.016.

(47) Amadeu, T. P.; Seabra, A. B.; de Oliveira, M. G.; Monte-Alto-Costa, A. Nitric Oxide Donor Improves Healing If Applied on Inflammatory and Proliferative Phase. Journal of Surgical Research 2008, 149 (1), 84–93. 10.1016/J.JSS.2007.10.015.

(48) Amadeu, T. P.; Seabra, A. B.; de Oliveira, M. G.; Costa, A. M. A. S-Nitrosoglutathione-Containing Hydrogel Accelerates Rat Cutaneous Wound Repair. Journal of the European Academy of Dermatology and Venereology 2007, 21 (5), 629–637. 10.1111/J.1468-3083.2006.02032.X.

(49) Lupu, A.; Bercea, M.; Avadanei, M.; Gradinaru, L. M.; Nita, L. E.; Gradinaru, V. R. Temperature Sensitive Pluronic F127-Based Gels Incorporating Natural Therapeutic Agents. Macromolecular Materials and Engineering 2025, 310 (4), 2400341. 10.1002/MAME.202400341.

(50) Li, J.; Li, X.; Zhou, Z.; Ni, X.; Leong, K. W. Formation of Supramolecular Hydrogels Induced by Inclusion Complexation between Pluronics and α-Cyclodextrin. Macromolecules 2001, 34 (21), 7236–7237. 10.1021/ma010742s.

(51) Wang, J.; Fang, Q.; Ye, L.; Zhang, A.; Feng, Z. G. The Intrinsic Microstructure of Supramolecular Hydrogels Derived from α-Cyclodextrin and Pluronic F127: Nanosheet Building Blocks and Hierarchically Self-Assembled Structures. Soft Matter 2020, 16 (25), 5906–5909. 10.1039/D0SM00979B.

(52) Polaschegg, H. D. Catheter Locking Solution Spillage: Theory and Experimental Verification. Blood Purification 2008, 26 (3), 255–260. 10.1159/000123706.

(53) Ash, S. R.; Mankus, R. A.; Sutton, J. M.; Criswell, R. E.; Crull, C. C.; Velasquez, K. A.; Smeltzer, B. D.; Ing, T. S. Concentrated Sodium Citrate (23%) for Catheter Lock. Hemodialysis International 2000, 4 (1), 22–31. 10.1111/HDI.2000.4.1.22.

(54) Lok, C. E.; Huber, T. S.; Lee, T.; Shenoy, S.; Yevzlin, A. S.; Abreo, K.; Allon, M.; Asif, A.; Astor, B. C.; Glickman, M. H.; Graham, J.; Moist, L. M.; Rajan, D. K.; Roberts, C.; Vachharajani, T. J.; Valentini, R. P. KDOQI Clinical Practice Guideline for Vascular Access: 2019 Update. American Journal of Kidney Diseases 2020, 75 (4), S1–S164. 10.1053/j.ajkd.2019.12.001.

(55) Akash, M. S. H.; Rehman, K. Recent Progress in Biomedical Applications of Pluronic (PF127): Pharmaceutical Perspectives. Journal of Controlled Release 2015, 209, 120–138. 10.1016/J.JCONREL.2015.04.032.

(56) Broniowska, K. A.; Diers, A. R.; Hogg, N. S-Nitrosoglutathione. Biochimica et Biophysica Acta (BBA)-General Subjects 2013, 1830 (5), 3173–3181. 10.1016/J.BBAGEN.2013.02.004.

(57) Avery, R. K.; Albadawi, H.; Akbari, M.; Zhang, Y. S.; Duggan, M. J.; Sahani, D. V.; Olsen, B. D.; Khademhosseini, A.; Oklu, R. An Injectable Shear-Thinning Biomaterial for Endovascular Embolization. Science Translational Medicine 2016, 8 (365). 10.1126/scitranslmed.aah5533.

(58) Zhao, Q.; Zhang, J.; Song, L.; Ji, Q.; Yao, Y.; Cui, Y.; Shen, J.; Wang, P. G.; Kong, D. Polysaccharide-Based Biomaterials with on-Demand Nitric Oxide Releasing Property Regulated by Enzyme Catalysis. Biomaterials 2013, 34 (33), 8450–8458. 10.1016/J.BIOMATERIALS.2013.07.045.

(59) Bohl, K. S.; West, J. L. Nitric Oxide-Generating Polymers Reduce Platelet Adhesion and Smooth Muscle Cell Proliferation. Biomaterials 2000, 21 (22), 2273–2278. 10.1016/S0142-9612(00)00153-8.

(60) Lipke, E. A.; West, J. L. Localized Delivery of Nitric Oxide from Hydrogels Inhibits Neointima Formation in a Rat Carotid Balloon Injury Model. Acta Biomaterialia 2005, 1 (6), 597–606. 10.1016/J.ACTBIO.2005.07.010.

(61) Taite, L. J.; West, J. L. Sustained Delivery of Nitric Oxide from Poly(Ethylene Glycol) Hydrogels Enhances Endothelialization in a Rat Carotid Balloon Injury Model. Cardiovascular Engineering and Technology 2011, 2 (2), 113–123. 10.1007/s13239-011-0040-z.

(62) Tabish, T. A.; Thorat, N. D.; Narayan, R. J. Mechanical Behaviour of Nitric Oxide Releasing Polymers for Cardiovascular Bypass Grafts. Mechanics of Materials 2023, 176, 104520. 10.1016/J.MECHMAT.2022.104520.

(63) Locatelli, F.; Buoncristiani, U.; Canaud, B.; Köhler, H.; Petitclerc, T.; Zucchelli, P. Dialysis Dose and Frequency. Nephrology Dialysis Transplantation 2005, 20 (2), 285–296. 10.1093/NDT/GFH550.

(64) Lok, C. E.; Mokrzycki, M. H. Prevention and Management of Catheter-Related Infection in Hemodialysis Patients. Kidney International 2011, 79 (6), 587–598. 10.1038/KI.2010.471.

(65) National Kidney Foundation. Catheter-Related Bloodstream Infection (CRBSI). 2024. https://www.kidney.org/kidney-topics/catheter-related-bloodstream-infection-crbsi

(66) Munoz-Mozas, G. Preventing Intravenous Catheter-Related Bloodstream Infections (CRBSIs). British Journal of Nursing 2023, 32 (Sup7), S4–S10. 10.12968/BJON.2023.32.SUP7.S4.

(67) Doellman, D. Guarding the Central Venous Access Device: A New Solution for an Old Problem. British Journal of Nursing 2023, 32 (19), S20–S25. 10.12968/BJON.2023.32.19.S20.

(68) Doverspike, J. C.; Zhou, Y.; Wu, J.; Tan, X.; Xi, C.; Meyerhoff, M. E. Nitric Oxide Releasing Two-Part Creams Containing S-Nitrosoglutathione and Zinc Oxide for Potential Topical Antimicrobial Applications. Nitric Oxide 2019, 90, 1–9. 10.1016/J.NIOX.2019.05.009.

(69) Doverspike, J. C.; Mack, S. J.; Luo, A.; Stringer, B.; Reno, S.; Cornell, M. S.; Rojas-Pena, A.; Wu, J.; Xi, C.; Yevzlin, A.; Meyerhoff, M. E. Nitric Oxide-Releasing Insert for Disinfecting the Hub Region of Tunnel Dialysis Catheters. ACS Applied Materials Interfaces 2020, 12 (40), 44475–44484. 10.1021/acsami.0c13230.

(70) Wang, D. C.; Clark, J. R.; Lee, R.; Nelson, A. H.; Maresso, A. W.; Acharya, G.; Shin, C. S. Development of Antimicrobial Nitric Oxide-Releasing Fibers. Pharmaceutics 2021, 13 (9), 1445. 10.3390/PHARMACEUTICS13091445.

(71) Yang, L.; Feura, E. S.; Ahonen, M. J. R.; Schoenfisch, M. H. Nitric Oxide–Releasing Macromolecular Scaffolds for Antibacterial Applications. Advanced Healthcare Materials 2018, 7 (13), 1800155. 10.1002/adhm.201800155.

(72) Arora, D. P.; Hossain, S.; Xu, Y.; Boon, E. M. Nitric Oxide Regulation of Bacterial Biofilms. Biochemistry 2015, 54 (24), 3717–3728. 10.1021/bi501476n.

(73) CDC. Guidelines for the Prevention of Intravascular Catheter-Related Infections. 2011. https://www.cdc.gov/infection-control/hcp/intravascular-catheter-related-infection/index.html

