## Supplementary Information for "Nitric Oxide-Releasing Thixotropic Hydrogels as Antibacterial and Hemocompatible Catheter Locks"

|  | 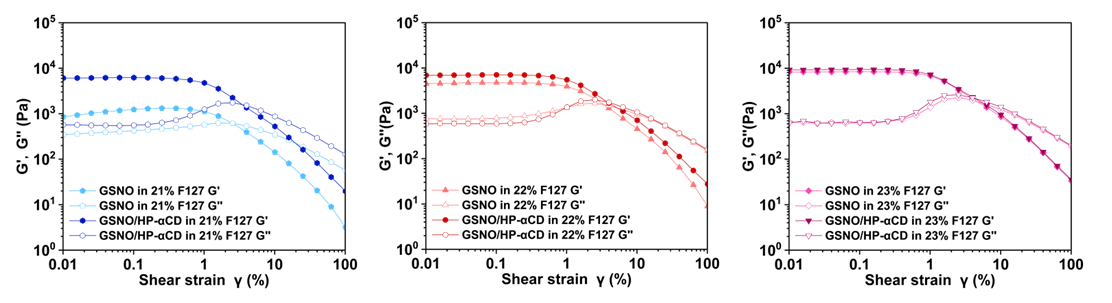 |
| --- | --- |
| Figure S1. Storage modulus (G′) and loss modulus (G'') of F127 hydrogels as a function of the strain amplitude at a frequency of 10 rad/s at 37 °C. | |

**Table S1.** Loss factor (tan δ) of various F127-based hydrogel formulations measured in amplitude sweep tests

| **Shear strain γ %** | **21%F127** | | **22% F127** | | |
| --- | --- | --- | --- | --- | --- |
|  | **GSNO** | **GSNO/HP-αCD** | **/** | **GSNO** | **GSNO/HP-αCD** |
| **0.0101** | 0.403 | 0.094 | 0.099 | 0.171 | 0.085 |
| **0.0159** | 0.387 | 0.093 | 0.092 | 0.167 | 0.086 |
| **0.0252** | 0.367 | 0.09 | 0.089 | 0.161 | 0.086 |
| **0.04** | 0.353 | 0.09 | 0.091 | 0.158 | 0.085 |
| **0.0634** | 0.344 | 0.088 | 0.088 | 0.162 | 0.085 |
| **0.101** | 0.341 | 0.09 | 0.09 | 0.165 | 0.084 |
| **0.159** | 0.343 | 0.094 | 0.092 | 0.171 | 0.086 |
| **0.252** | 0.352 | 0.105 | 0.097 | 0.185 | 0.09 |
| **0.4** | 0.373 | 0.123 | 0.13 | 0.212 | 0.108 |
| **0.634** | 0.414 | 0.167 | 0.154 | 0.255 | 0.148 |
| **1.01** | 0.51 | 0.265 | 0.261 | 0.342 | 0.248 |
| **1.59** | 0.709 | 0.471 | 0.481 | 0.522 | 0.446 |
| **2.53** | ***1.001*** | 0.779 | 0.769 | 0.798 | 0.744 |
| **4** | 1.363 | ***1.119*** | ***1.022*** | ***1.142*** | ***1.043*** |
| **6.34** | 1.823 | 1.374 | 1.229 | 1.572 | 1.229 |
| **10.1** | 2.437 | 1.66 | 1.547 | 2.11 | 1.498 |
| **15.9** | 3.267 | 2.085 | 1.969 | 2.774 | 1.911 |
| **25.2** | 4.453 | 2.688 | 2.54 | 3.7 | 2.468 |
| **40** | 6.286 | 3.567 | 3.418 | 5.398 | 3.265 |
| **63.4** | 9.628 | 4.786 | 4.637 | 8.651 | 4.371 |
| **101** | 17.841 | 6.474 | 6.125 | 16.743 | 5.732 |

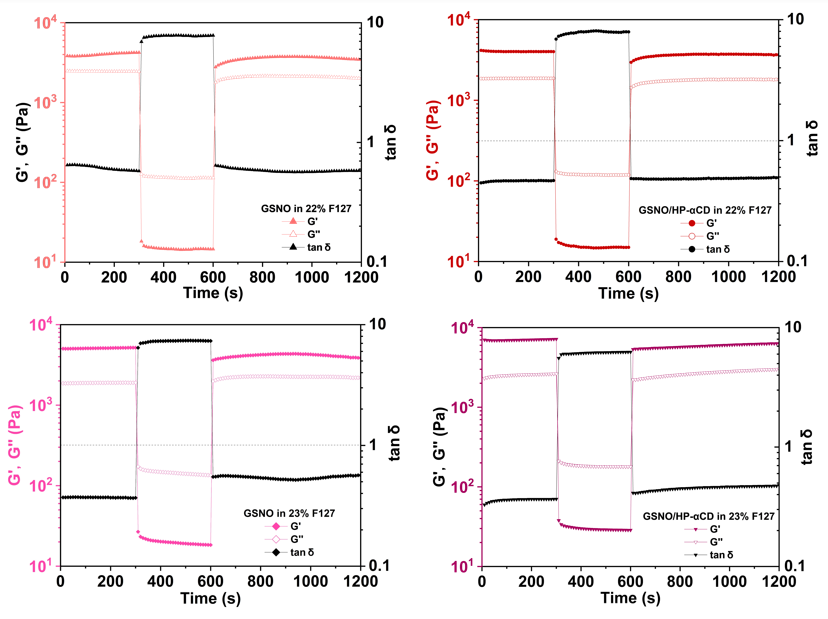

Figure S2. Storage modulus (G′), loss modulus (G''), and loss factor (tan δ) of various F127 hydrogels as a function of time during successive step strain measurements at low (1%), high (100%), and low (1%) shear strains at 37°C.

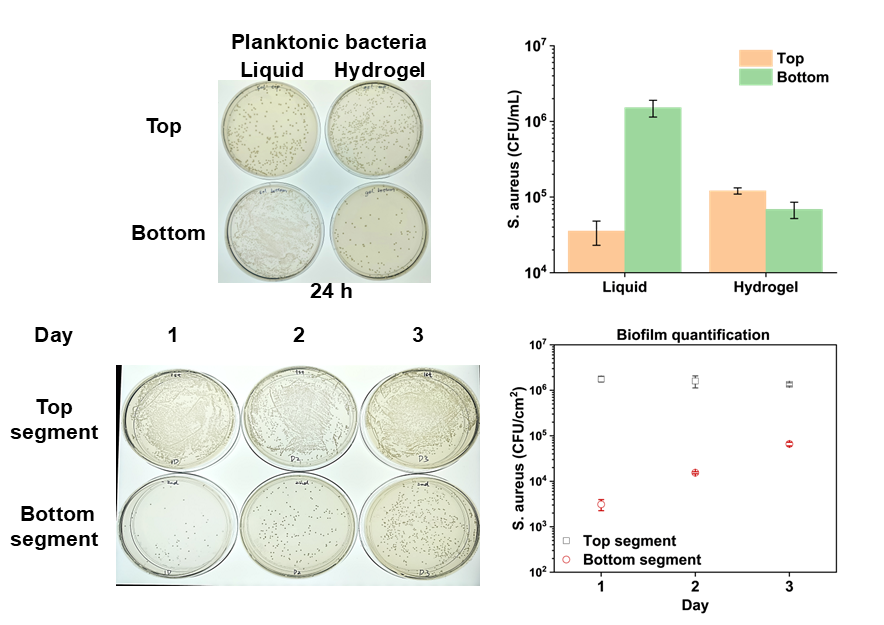

Figure S3. Quantification of *S. aureus* biofilm in the top and bottom segments of the catheter filled with a drug-free F127 hydrogel.

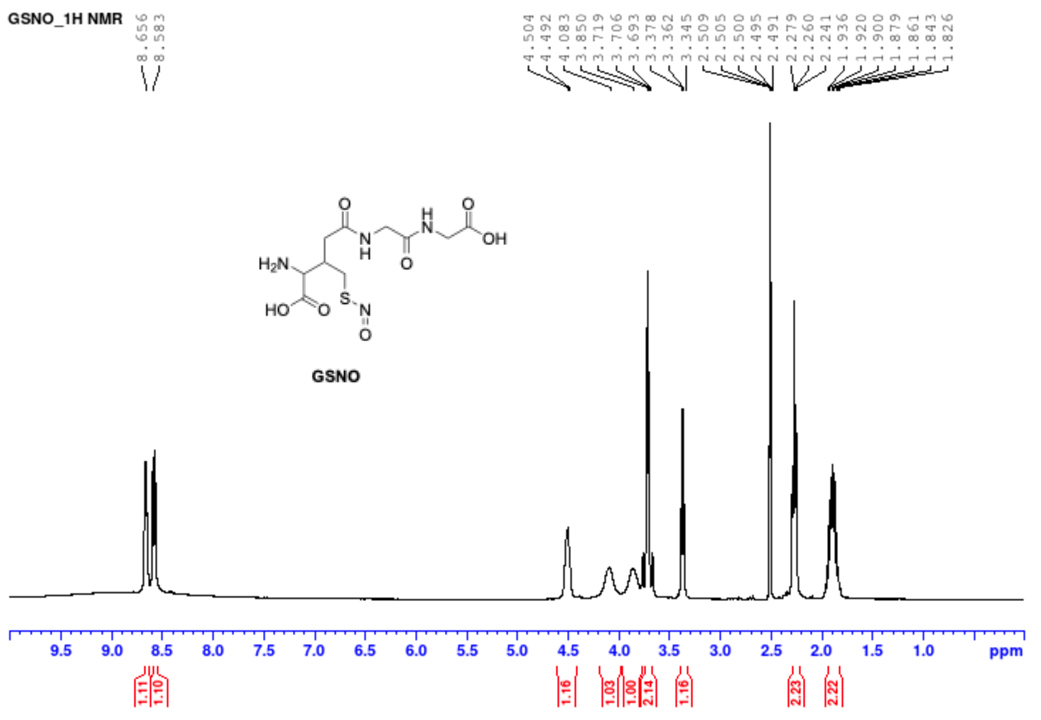

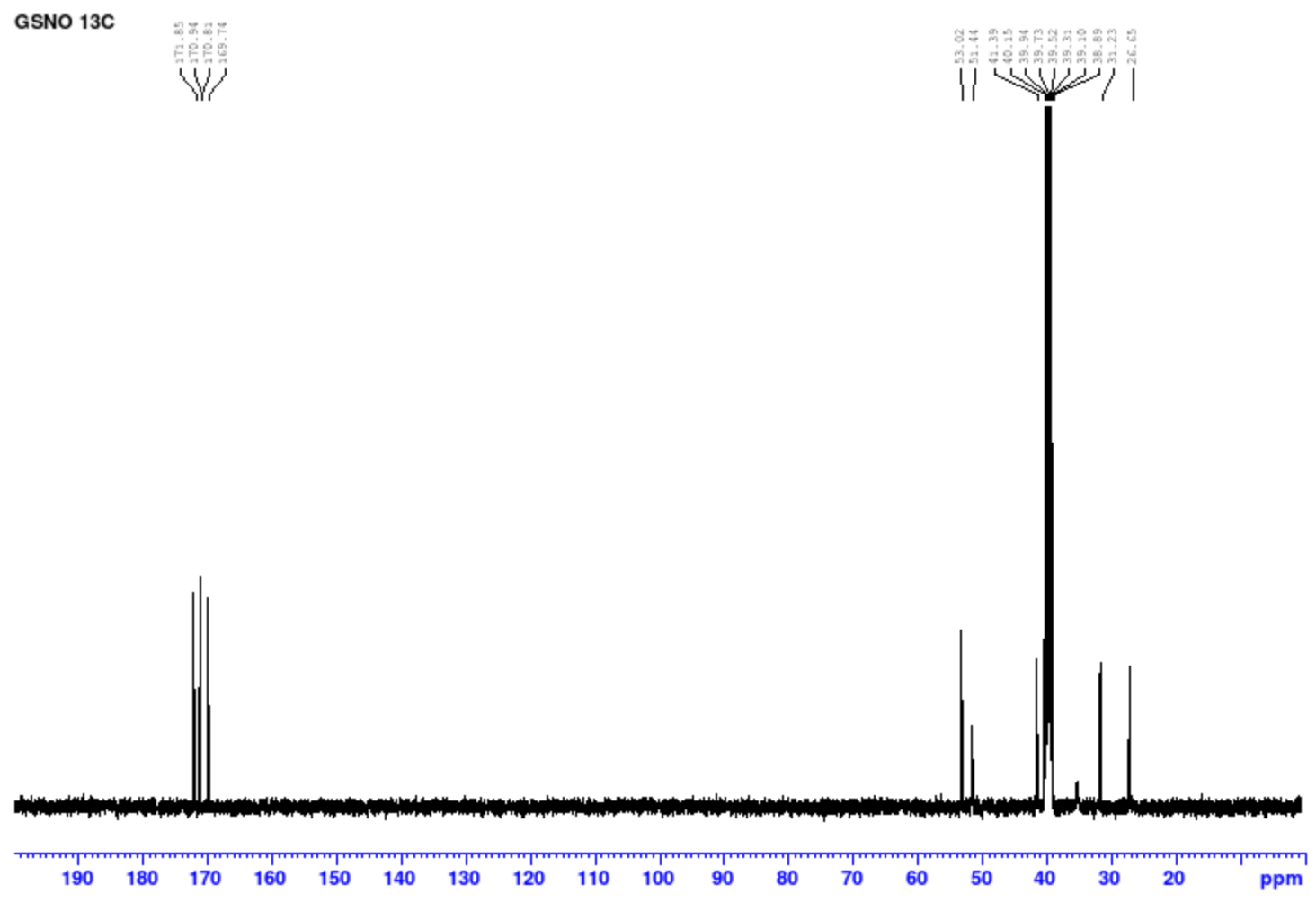

Figure S4. ^1^H NMR spectrum and ^13^C NMR spectrum of GSNO.

^1^H NMR (400 MHz, DMSO): δ 8.65 (bs, 1H), 8.57 (bs, 1H), 4.50 (d, J = 5Hz, 1H), 4.15 ~ 4.0 (m, 1H), 3.92 ~ 3.80 (m, 1H), 3.71 ~ 3.69 (m, 2H), 3.36 (t, J = 6.35, 12.7 Hz, 1H), 2.25 (t, J = 7.44, 14.94 Hz, 2H), 1.93-1.82 (m, 2H).

^13^C NMR (100 MHz, DMSO): δ; 171.8, 170.9, 170.8, 169.7, 53.0, 51.4, 41.3, 39.9, 31.2, 26.6.
